# Effect of Microbubble Size, Composition and Multiple Sonication Points on Sterile Inflammatory Response in Focused Ultrasound-Mediated Blood-Brain Barrier Opening

**DOI:** 10.1101/2024.04.28.591538

**Authors:** Payton J. Martinez, Jane J. Song, Jair Castillo, John DeSisto, Kang-Ho Song, Adam L. Green, Mark Borden

## Abstract

Blood-brain barrier opening (BBBO) using focused ultrasound (FUS) and microbubble (MBs) has emerged as a promising technique for delivering therapeutics to the brain. However, the influence of various FUS and MB parameters on BBBO and subsequent sterile inflammatory response (SIR) remains unclear. In this study, we investigated the effects of MB size and composition, as well as the number of FUS sonication points, on BBBO and SIR in an immunocompetent mouse model. Using MRI-guided MB+FUS, we targeted the striatum and assessed extravasation of an MRI contrast agent to assess BBBO and RNAseq to assess SIR. Our results revealed distinct effects of these parameters on BBBO and SIR. Specifically, at a matched microbubble volume dose (MVD), MB size did not affect the extent of BBBO, but smaller (1 µm diameter) MBs exhibited a lower classification of SIR than larger (3 or 5 µm diameter) MBs. Lipid-shelled microbubbles exhibited greater BBBO and a more pronounced SIR compared to albumin-shelled microbubbles, likely owing to the latter’s poor *in vivo* stability. As expected, increasing the number of sonication points resulted in greater BBBO and SIR. Furthermore, correlation analysis revealed strong associations between passive cavitation detection measurements of harmonic and inertial MB echoes, BBBO and the expression of SIR gene sets. Our findings highlight the critical role of MB and FUS parameters in modulating BBBO and subsequent SIR in the brain. These insights inform the development of targeted drug delivery strategies and the mitigation of adverse inflammatory reactions in neurological disorders.

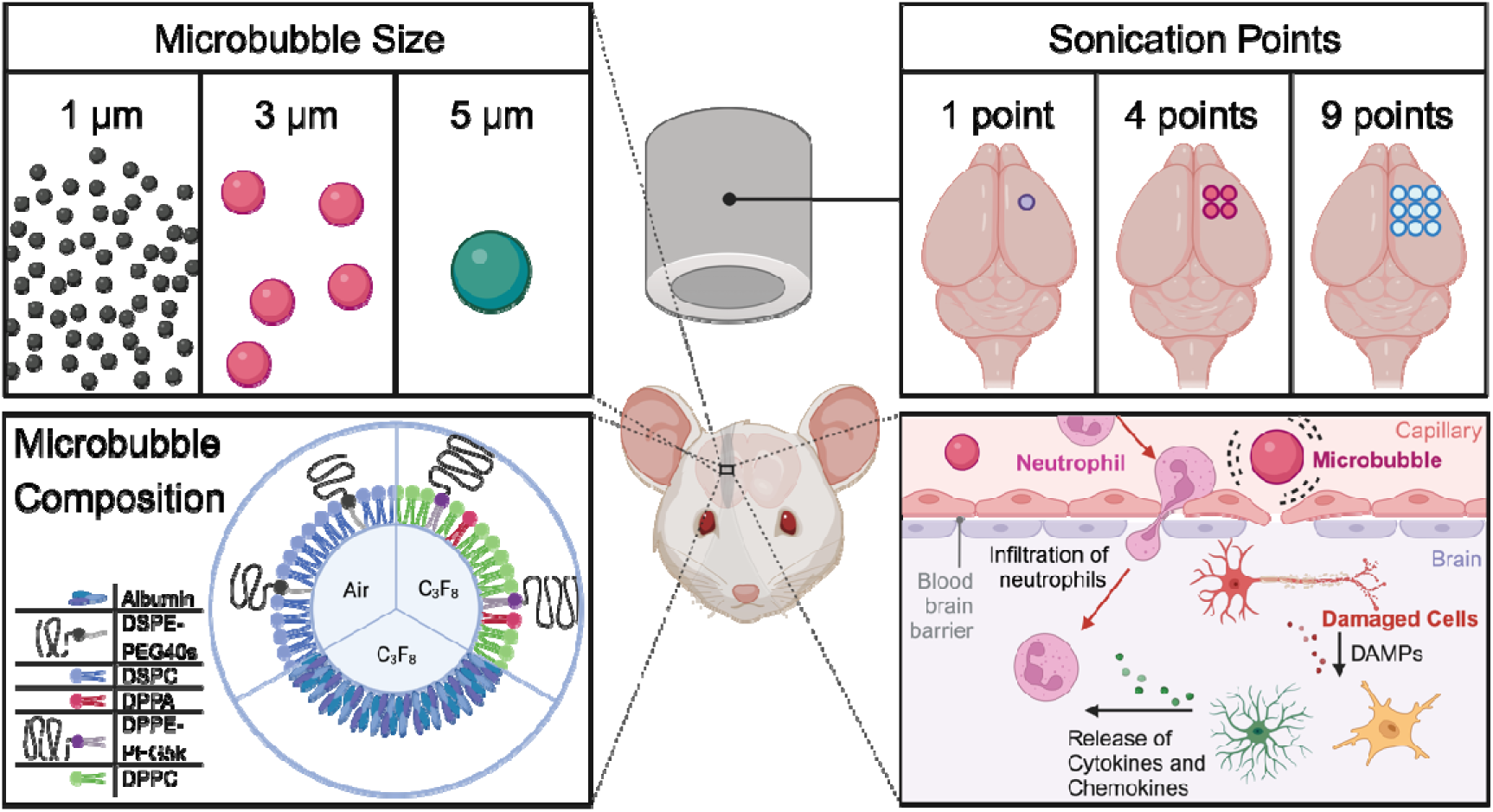

## INTRODUCTION

The blood-brain barrier (BBB) is a critical biological barrier influencing the delivery of drugs and imaging agents into the brain parenchyma. In 2001, Hynynen et al. performed the first blood-brain barrier opening (BBBO) experiment using focused ultrasound (FUS) and systematically circulating microbubbles (MBs).^1^ This technique has been further developed to deliver drugs including chemotherapeutics,^2–4^ liposomes,^5^ antibodies^6^ and other therapeutic molecules past the BBB. Using clinically approved MBs, clinical trials have expanded to include multiple pathologies (e.g., glioblastoma, diffuse midline gliomas, brain metastasis, amyotrophic lateral sclerosis, Parkinson’s disease and Alzheimer’s disease).^7–9^

A microbubble is a 1-10 µm diameter colloidal particle with a thin viscoelastic shell (e.g., protein, lipid or polymer) that encapsulates a high-molecular-weight, low-solubility gas (e.g. perfluorobutane, sulfur hexafluoride) core, suspended in an aqueous medium.^10^ MBs were first developed as ultrasound contrast agents and clinically approved for echocardiography and other imaging applications in the late 1990s. For imaging, clinically approved MB suspensions are polydisperse in size, comprise 10^8^ to 10^9^ MBs/mL and are typically injected intravenously to a microbubble volume dose (MVD) of 0.01-10 µL/kg (Table S1). Higher MVDs (10-40 µL/kg) are typically used for MB+FUS applications.^11^ Pharmacokinetics determine the number and size of MBs in the local brain microvasculature of the FUS focal region at a given time. For a given MB composition, pharmacokinetics were found to be primarily affected by MVD.^12^ MB composition, for example the lipid species or concentration of poly(ethylene glycol) (PEG) on the lipid shell, can also influence pharmacokinetics.^13,14^

When exposed to an acoustic field, the rarefaction and compression within the ultrasound focus cause volumetric oscillations due to its compressible gas core, providing the localized forces needed to disrupt the BBB. The behavior of MBs under ultrasound can vary significantly. At lower pressures, they stably oscillate harmonically; whereas at higher pressures they can expand to a point where their collapse causes the MB to collapse inertially and emit an acoustic shock wave.^15–17^ Both regimes of cavitation generate mechanical forces, such as contact, shear and reradiated acoustic forces, that impinge on the endothelium and other elements of the BBB.^11,18^ Tight junctions, which regulate the porosity of the vessel lumen, react to these forces by mechanically disassembling, creating gaps along the BBB.^19^ However, excessive MB oscillation can damage the BBB and nearby cells beyond what is necessary for drug delivery. Different MB acoustic echoes emerge from these regimes: small oscillations produce harmonic frequencies (subharmonics, fundamental and ultra-harmonics), whereas strong oscillations with inertial collapse generate shockwaves detected by their broadband frequency content.^15,16^ Real-time insights into MB activity within the FUS beam can be obtained using passive cavitation detection (PCD).

Disruption of the BBB may alter the biological microenvironment within the ultrasound focal region. Previous studies have reported BBBO-induced microhemorrhages, transient edema and even cell death.^20^ With advancements in transcriptomics, more sophisticated examinations of BBBO effects have shown upregulation in major inflammatory pathways, including the NFκB,^21,22^ Jak-Stat^23^ and apoptosis^23^ pathways. This inflammatory response, occurring without infection, mirrors the sterile inflammatory response (SIR) seen in other neurological injuries (e.g. traumatic brain injury, stroke), initiated by damage-associated molecular patterns (DAMPs) released from injured cells.^24–26^ Nearby immune cells (astrocytes and microglia) recognize and amplify these signals, leading to an inflammatory peak between 3 and 12 hours.^21,27^ Although many studies have shown windows of this response, the complete mechanism and influence of MB and FUS parameters is still unclear.^28–32^

The uniquely modular nature of MBs and FUS has allowed the manipulation of parameters to optimize drug delivery. Over the past two decades, much research has gone into improving these parameters, including pulse length,^33,34^ pulse repetition frequency^33,35,36^ and sonication duration.^37^ Recent developments have introduced the concept of multiple sonication points to increase the size of the treated region, although their effect beyond increasing BBBO volume has yet to be fully explored. Likewise, microbubble parameters including size, composition and dose have received less attention, possibly due to the reliance on clinically approved, polydisperse agents.^38,39^ In 2017, a debate between Kovacs et al.^40,41^ and McMahon et al.^42,43^ elucidated the importance of these parameters on the SIR.

In this study, we aim to unravel the importance of MB properties (size and composition), especially in the context of multiple sonication points (Figure 1). Utilizing MRI-guided MB+FUS, we targeted the striatum to deliver an MRI intravascular contrast agent (gadobenate dimeglumine) to quantify the extent of BBBO. Samples from the affected and contralateral regions were collected for bulk RNA sequencing 6 h post-MB+FUS. By varying the number of sonication points (1, 4 or 9), microbubble size (1, 3 and 5 µm diameter) and microbubble composition, we illustrate the effect of each on both the extent of BBBO and its subsequent SIR. Finally, we correlate the results to determine the most important parameters in attaining BBBO while limiting SIR.

**Figure 1:**
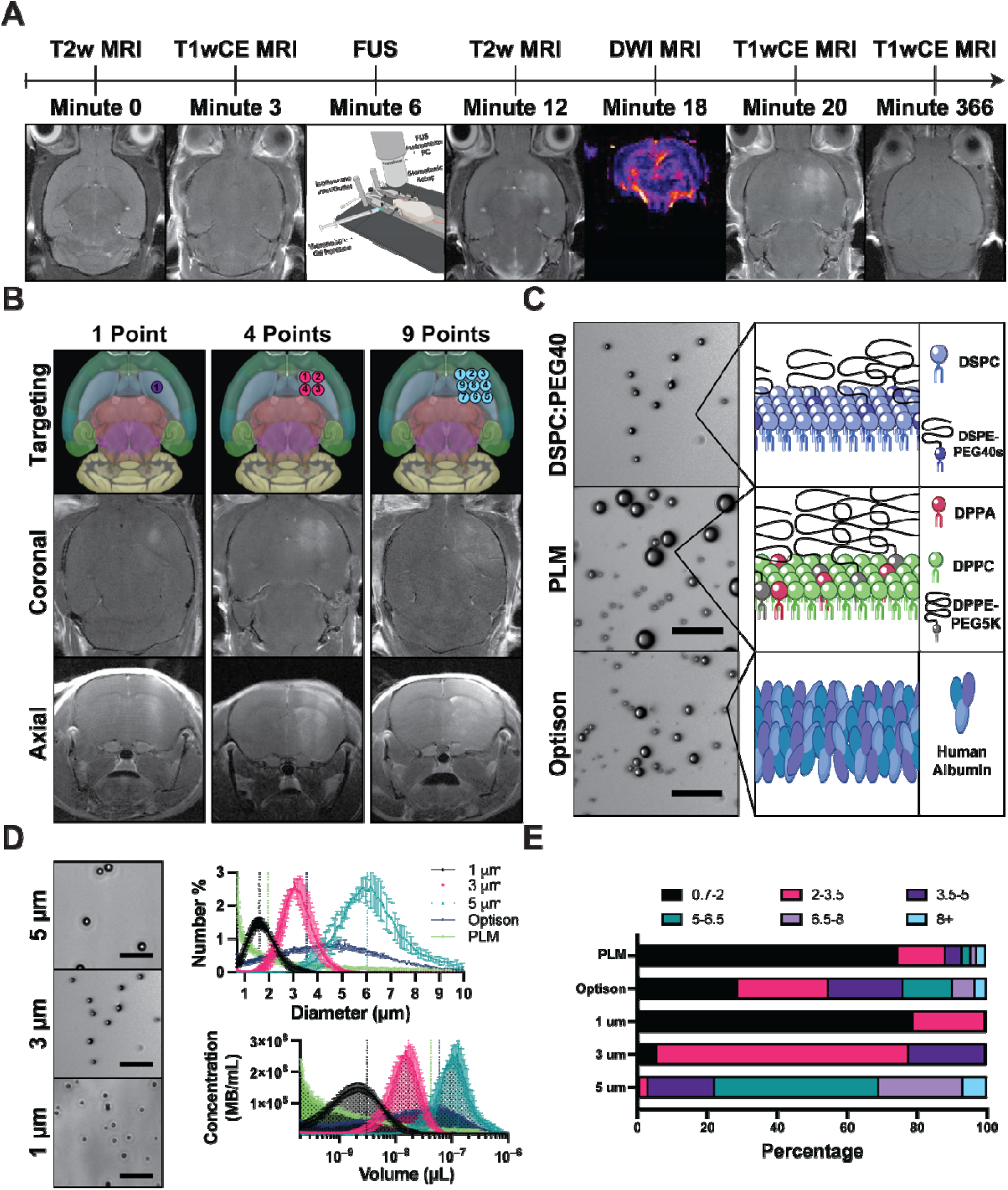
Experimental Set Up and Microbubble Characterization. (A) Timeline of FUS treatment. Initial MRI is done including T2w (targeting) and T1w (BBB integrity). Mice are stereotactically moved to the FUS system (RK50) and sonicated for 3 min. Mice are then moved back to MRI for post-MB+FUS imaging, which involves T2w (edema), DWI (hemorrhaging) and T1w (extent of BBBO). (B) Three targeting schemes were employed: 1 point (left), 4 points (middle) and 9 points (right), all centered on the striatum (light blue region). A representative post-MB+FUS T1w MRI coronal (middle) and axial (bottom) is also displayed. (C) Three microbubble compositions were tested: DSPC:PEG40S (top), PLM (middle) and Optison (bottom). Representative brightfield images of each microbubble sample are shown on the left, with a cartoon illustrating the shell composition on the right-hand side. (D) Microbubbles using the DSPC:PEG40S composition were isolated into three sizes (1, 3 and 5 μm diameter). Representative brightfield images ar displayed on the left, accompanied by size distributions showing the number-weighted diameter (top) and concentration-weighted volume (bottom) on the right. (E) A bar chart depicting the sizes of all tested bubbles, with each color representing a range of diameters. All data are presented as mean ± SD.

## MATERIALS AND METHODS

### Materials

All solutions were prepared using filtered, sterile, deionized water (Direct-Q 3 Millipore, Billerica, MA, USA. 1,2-distearoyl-sn-glycero-3-phosphocholine (DSPC), 1,2-dipalmitoyl-sn-glycero-3-phosphate (DPPA) and 1,2-dipalmitoyl-sn-glycero-3-phosphocholine (DPPC) were purchased from Avanti Polar Lipids (Alabaster, AL, USA). Perfluorobutane gas (n-C4F10, PFB) was purchased from FluroMed (Round Rock, TX, USA). Polyoxyethylene-40 stearate (PEG40S), 1,2-dipalmitoyl-sn-glycero-3-phosphoethanolamine-N-[methoxy(polyethylene glycol)-5000] (DPPE-PEG5K) and chloroform were purchased from Sigma-Aldrich (St. Louis, MO, USA). Phosphate-buffered saline was procured from Fisher Scientific (Pittsburg, PA, USA). All reagents were ≥99% pure and used as received without further purification.

### Microbubble Suspensions

Microbubbles comprising DSPC:PEG40S at 90:10 molar ratio were prepared as previously described by Fesitan et al.^44^ Under sterile conditions, polydisperse MBs were created by sonication of the lipid suspension with decafluorobutane gas and collected. Three diameters (1, 3 and 5 ± 0.5 µm) were isolated by differential centrifugation (Table 1). The isolation process, including centrifugation speeds used, can be found in supplemental Figure S1. The size-isolation process was performed in air-saturated media, and thus the resulting DSPC:PEG40S MBs likely comprised air prior to injection due to gas exchange.^45,46^ The size isolation method also “washes” the MBs of free lipid prior to injection.

**Table 1:**
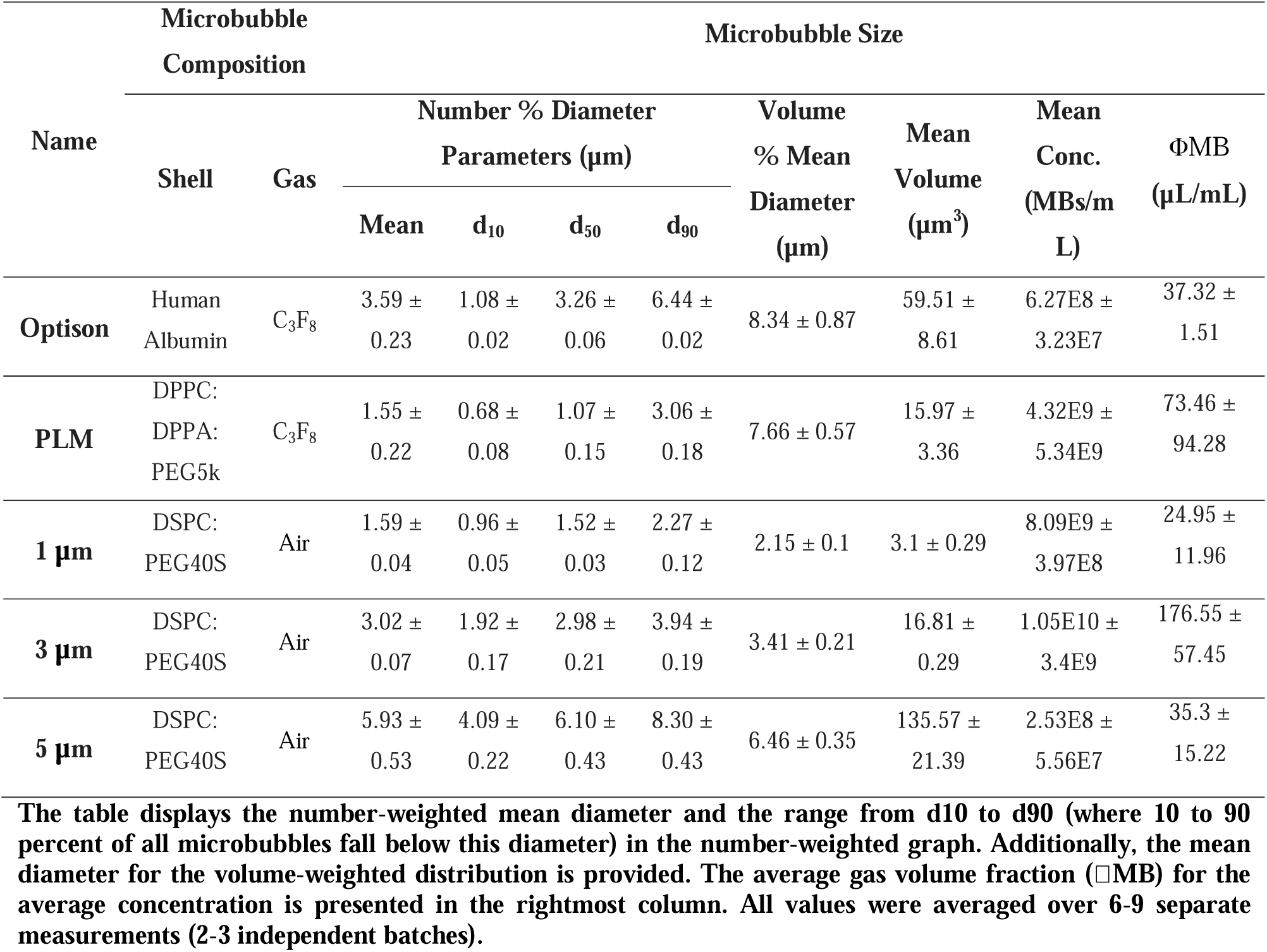
Microbubble formulation characterization.

“Definity-like” (perflutren lipid microspheres) MBs were created using a similar composition to the clinically approved ultrasound contrast agent Definity (Lantheus) (DPPC:DPPA:PEG5k at 82:10:8 molar ratio to 0.75 mg/mL total lipid concentration in 10 vol% glycerin, 10 vol% propylene glycol and 80 vol% PBS) with a headspace of octafluoropropane in a 2-mL serum vial capped and sealed (Table 1). MBs were formed by agitation (activation) for 45 s in a dental amalgamator to create perflutren lipid microspheres (PLMs). These MBs were stored immediately following activation and injected up to 6 hours later and therefore comprised an octafluoropropane core, and the suspension had free lipid in the form of liposomes and/or micelles.

Optison microbubbles were purchased directly from GE Healthcare and resuspended as directed on the package insert. These MBs comprised a human serum albumin protein shell and an octafluoropropane gas core. These MBs were injected immediately following activation and therefore comprised an octafluoropropane core, and the suspension had free protein from the manufacturing process and any destabilized MBs due to storage and handling.

Microbubble concentration and number- and volume-weighted size distributions were measured with a Multisizer 4 (Beckman Coulter). Microbubble concentration (*c_i_*, MBs/µL) versus microbubble volume (*v_i_*, µL/MB) was plotted, and the gas volume fraction (*ϕMB*) was estimated as follows:

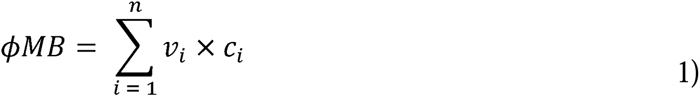

where *i* is the index of the sizing bin, with 300 bins ranging from 0.7 to 18 µm in diameter. Three independent MB preparations were measured two hours before FUS treatment to confirm size distributions and concentration. Microbubble cakes were stored in the refrigerator at 4 °C for use within 3 days. Microbubbles were diluted to achieve the desired microbubble volume dose (MVD) injection concentration within 2 min before injection.

### MRI-Guided MB+FUS BBBO Procedure

All animal experiments were conducted according to the regulations and policies of the University of Colorado Anschutz Institutional Animal Care and Use Committee (IACUC) in CD-1 IGS mice. All mice used were female, 8 to 11 weeks old, and purchased from Charles River Laboratory (Wilmington, MA).

A Bruker BioSpec 9.4/ 20 Tesla magnetic resonance imaging (MRI) Scanner (Bruker, Billerica, MA) with a mouse head RF phase-array coil was used in the Colorado Animal Imaging Shared Resource (RRID:SCR_021980). Mice were placed into a custom-designed MRI bed that contained stereotaxic ear bars to prevent movement of the mouse head during the transfer from MRI to the focused ultrasound (FUS) system and back into the MRI scanner. Each MRI session consisted of: (I) a 3D-localizer, a T1w MSME (Multi-Spin Multi-Echo) image performed 14 min after intravenous injection of 0.4 mmol/kg gadobenate dimeglumine (MultiHance, Bracco, Princeton, NJ) (II) high-resolution 3D T2-weighted turboRARE (Rapid Acquisition with Relaxation Enhancement, 52 µm in-plane resolution); and (III) axial fast spin echo DWI (Diffusion Weighted Imaging) with six b-values for tumor cellularity and edema.^47^ Mice remained stereo-tactically placed on an MRI bed and transferred to the FUS system for treatment where an intravenous injection of 0.1 mL gadobenate dimeglumine (MultiHance, Bracco, Milan, Italy) was given. All image acquisition was performed using Bruker ParaVision NEO360 v.3.3 software. A timeline of treatments can be found in Figure 1.

During the MB+FUS application, a single element, geometrically focused transducer (frequency: 1.0 MHz, diameter: 30 mm) was driven by the RK-50 system (FUS Instruments, Toronto, Canada). A single element, geometrically focused transducer (frequency: 0.5 MHz, diameter: 10 mm) coaxially inside the driving transducer was used for passive cavitation detection (PCD). The experimental setup is shown in Fig. 1A. Using the coronal T2-weighted MR image (coronal), the center of the striatum was targeted with either 1, 4 or 9 points (Fig. 1B). Ultrasound gel (Aquasonic gel, Clinton Township, MI) was placed on the mouse head confirming the lack of air bubbles. An acoustically transparent tank filled with degassed water was placed on top of the gel (Fig. 1D). Microbubbles (10 μL/kg; 0.1 mL) and 0.1 mL MultiHance were injected intravenously through a retroorbital injection via a 26 Ga needle. Directly after injection, FUS was applied. FUS parameters were as follows: 10 ms PL, 1 Hz PRF, 180 s treatment time, and a peak negative pressure (PNP) of 0.5 MPa (0.4 MPa in situ). Directly after MB+FUS, mice were then sent back to MRI to complete post-MB+FUS T1-weighted MRI. All groups were randomly selected (n = 3 mice/group).

### Passive Cavitation Detection Collection and Analysis

Voltage data from the PCD was collected from 0.5 ms before to 0.5 ms after sonication at each sonication point separately. Each sonication point and ultrasound burst was analyzed as previously described^48^ resulting in harmonic cavitation content (HCC) and broadband cavitation content (BCC) for each burst throughout the treatment. The remaining PCD analysis was done using MATLAB (Massachusetts, USA).

### MRI Analysis

T1w MRI datasets were used to quantify the extent of BBBO using FIJI software (Maryland, USA). All axial slices were analyzed by defining the contralateral hemisphere and determining the mean and standard deviation of voxel intensities. The treated hemisphere was then defined, and all voxels were found above two standard deviations of the contralateral side. The area was determined and multiplied by slice thickness (0.7 mm) to determine the BBBO volume. The contrast enhancement was determined by the average intensity within the BBB opening volume and divided by the average intensity of the control region. T2-weighted MRI was checked for edema and hemorrhaging by analyzing hyper and hypo-intense regions. Similar to T1w MRI, axial slices were analyzed by defining the contralateral hemisphere and determining the mean and standard deviation of voxel intensities. The treated hemisphere was then defined, and all voxels were found above two standard deviations of the contralateral side. Significant edema or hemorrhaging was defined by a region larger than one standard deviation above control mice (only given isoflurane, no FUS or MBs).

### Mouse Perfusion and Dissection

At 6 h post MB+FUS treatment, mice were sacrificed via perfusion with 25 mL of ice-cold PBS. Brains were immediately dissected using MRI guidance to only remove regions that were sonicated. Both the treated side and contralateral regions were removed and snap-frozen using liquid nitrogen. Samples were stored at −80 °C until further use. Brain samples were weighed and then immediately placed into a cell lysing buffer (Qiagen, Hilden, Germany), and homogenized for 30 s. RNA was isolated and purified using the RNAeasy Kit (74004, Qiagen), following the reagent manufacturer’s instructions. Quality control and library preparation were performed through the Anschutz Genomics Core for sequencing. Poly A selected total RNA paired-end sequencing was conducted at 40 million paired reads (80 million total reads) on a NovaSEQ 6000 sequencer.

Another subset of mice underwent histological analysis and were treated/perfused as described above. However, rather than a dissection of both sides, the brain was sliced axially at the center of the treated region and was immediately put into a 10% formalin solution and left to shake on an orbital shaker overnight at room temperature.

### Bulk RNA Sequencing Analysis

FASTQ files were obtained from the University of Colorado Anschutz Genomics Core after sequencing. RNA Analysis was performed using Pluto software (https://pluto.bio). Differential expression analysis was performed by comparing experimental groups to a control group (+Isoflurane). Genes were filtered to include only genes with at least 3 reads counted in at least 20% of samples in any group. Differential expression analysis was then performed with the DESeq2 R package,^49^ which tests for differential expression based on a model using the negative binomial distribution. Log2 fold change was calculated for the comparison of the experiment to the control group. Thus, genes with a positive log2 fold change value were deemed to have increased expression in the samples. Genes with a negative log2 fold change value had increased expression in control samples. Gene set enrichment analysis (GSEA) was performed using the fgsea R package and the fgseaMultilevel function.^50^ The log2 fold change from the experiment vs. control differential expression comparison was used to rank genes. Hallmarks gene set collection from the Molecular Signatures Database (MSigDB)^51,52^ was curated using the msigdbr R package.

### Statistical Analysis

All data are presented as mean ± SD. No preprocessing was done to the data except for voltage data collected from the PCD. PCD data were preprocessed as described in Martinez et al.^48^ All statistical analysis was completed in Prism 9 (GraphPad, California, USA). Star representations of p-values are indicated in captions and less than 0.05 was indicative of statistical significance. An unpaired Students’ t-test and ANOVA/multiple comparisons were used to compare two groups and larger comparisons, respectively. The false discovery rate (FDR) method was applied for multiple testing corrections.^53^ An adjusted p-value of 0.01 was used as the threshold for statistical significance.

## RESULTS AND DISCUSSION

### Characterization of Microbubble Properties Reveals Differences in Size and Composition

Three MB compositions were characterized prior to use in MB+FUS experiments. The first formulation, which has been previously characterized and used in MB+FUS,^12,48^ was DSPC:PEG40S-coated and air-filled MBs. Figure 1C shows a brightfield image of these MBs at an isolated size of 3-µm diameter. Next, we used a composition and creation method similar to the clinically approved Definity (PLM) (Fig. 1C). Lastly, protein-shelled Optison MBs were directly sourced from GE Healthcare and prepared according to the package insert instructions. Under brightfield microscopy, both PLM and Optison MBs displayed polydisperse populations (Fig. 1C).

To investigate the effect of MB size, the DSPC:PEG40S microbubbles were size isolated into three diameters (1, 3 and 5 µm). Brightfield microscopy confirmed the monodisperse nature of these MB populations (Fig. 1D). The number-weighted mean diameters for the tested microbubble populations in ascending order were as follows: PLM (1.55 ± 0.22 µm), 1 µm (1.59 ± 0.04 µm), 3 µm (3.02 ± 0.07 µm), Optison (3.59 ± 0.23 µm), and 5 µm (5.93 ± 0.53 µm). The volume-weighted mean diameters in ascending order were found to be PLM (7.66 ± 0.57 µm), 1 µm (2.15 ± 0.10 µm), 3 µm (3.41 ± 0.21 µm), 5 µm (6.46 ± 0.35 µm) and Optison (8.34 ± 0.27 µm), and. The volume-weighted size distribution plots are shown in Supplemental Figure S2. Other parameters, including gas volume fraction and concentration, can be found in Table 1. To accurately obtain a 10 µL/kg MVD, each microbubble preparation was measured and analyzed to find the gas volume fraction (*ϕMB*). The resulting values were: 73.5 ± 94.3, 37.3 ± 1.5, 25.0 ± 12.0, 176.6 ± 57.5, 35.3 ± 15.2 µL/mL for PLM, Optison, 1 µm, 3 µm and 5 µm MBs, respectively. A comparison of the five populations based on size highlighted that PLMs most closely resembled the 1 µm MBs. Due to its high polydispersity, the average number-weighted size of Optison was most similar to the 3 µm MBs and the volume-weighted size was most similar to 5 µm (Fig. 1E).

Currently, there is a wide range of ultrasound contrast agents used for MB+FUS BBBO.^7–9^ However, all of these polydisperse agents were originally formulated for echocardiographic contrast-enhanced ultrasound imaging. The variability in concentration and mean volume between batches, as illustrated in the gas volume fraction (Fig, 1D and Table 1), suggests that using a constant fluid volume (mL/kg) may yield inconsistent results. In this study, all formulations were matched to the desired MVD of 10 µL/kg before injection to mitigate such variations.

### PCD Shows Little Variation with Microbubble Size, Composition and Number of Sonication Points

Passive cavitation detection (PCD) was used to monitor MB activity throughout each MB+FUS sonication (Fig. 2A). The total harmonic cavitation content (HCC) was found for all groups (Fig. 2B), illustrating the increased total cavitation activity when a larger number of sonications were used. At the same number of points (4), protein-shelled Optison had significantly less HCC compared to the lipid-shelled DSPC:PEG40S MBs. We did not observe a significant difference between PLM and DSPC:PEG40S or between the differently sized DSPC:PEG40S formulations at this matched MVD (10 µL/kg).

**Figure 2:**
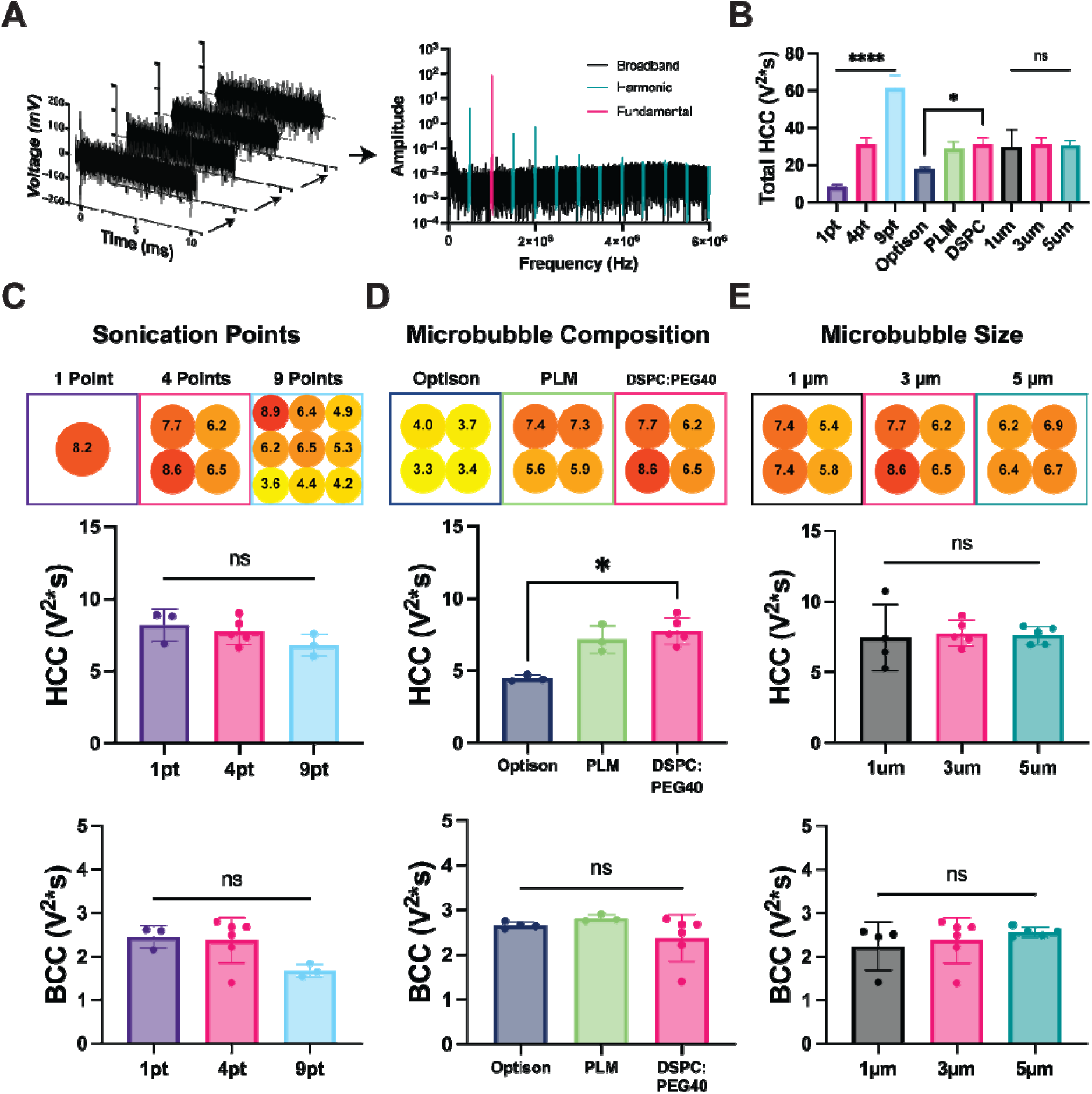
Quantification of Cavitation Content via Passive Cavitation Detection. (A) Schematic illustratin the method for quantifying PCD signal intensity into two cavitation contents. (B) Bar plot comparing the total harmonic cavitation content (HCC) values for all groups. Average HCC/BCC (broadband cavitation content) per point was quantified for each group and compared between variations in sonication points (C), composition (D) and size (E). Each panel (C-E) displays the harmonic cavitation content for each sonication point individually (top). The average harmonic cavitation content for all points (middle) and the average broadband cavitation content for all points (bottom) are also depicted. Data for 4 points / 3 µm / DSPC:PEG40S was repeated to facilitate direct comparisons with other groups. All data are presented as mean ± SD. * p<0.05, **** p<0.001.

To remove the influence of sonication points, each treatment was therefore averaged for all points. Figure 2C shows the changes in MB activity between the three sonication methods. The 1- and 4-point sonications displayed consistent levels of activity, whereas the 9-point sonication exhibited a bias towards the upper left region. This discrepancy is likely due to the skull’s shape in this area, being more perpendicular in the upper left and starting to curve at the lower right side (Supplemental Fig. S3). Upon investigation into average HCC for all points, we found similar values with a slight numerical decrease as the number of points increased (8.2 ± 1.1, 7.4 ± 1.3 and 6.8 ± 0.8 V2*s for 1, 4 and 9 points respectively) (Fig. 2C). No significant differences in broadband cavitation content (BCC) were noted, with mean values as follows: 2.5 ± 0.3, 2.4 ± 0.5 and 1.7 ± 0.2 V2*s for 1, 4 and 9 points respectively (Fig. 2C).

Microbubble composition showed consistency in HCC between all sonication targets (Fig. 2D, top). However, further analysis of average HCC revealed differences between groups, particularly with Optison MBs (4.5 ± 0.2 V2*s) showing overall lower activity than PLM (7.2 ± 1.0 V2*s) or DSPC:PEG40S (7.4 ± 1.3 V2*s) MBs (Fig. 2D, middle). These differences were not translated to inertial BCC, where there were no significant differences between groups (2.7 ± 0.1, 2.8 ± 0.1 and 2.4 ± 0.5 V2*s for Optison, PLM and DSPC:PEG40S MBs, respectively) (Fig. 2D, bottom).

In this study, using constant ultrasound PNP and microbubble MVD, we observed similar HCC across all groups with one exception: Optison MBs demonstrated significantly lower HCC compared to PLM or DSPC:PEG40S MBs (Fig. 2D, middle). Other researchers have reported similar results,^54–56^ although we have not directly compared their pharmacokinetics in vivo at a matched MVD. Bing et al. illustrated a 3-fold increase in the total area-under-the-curve (AUC) of passive cavitation feedback from Definity to Optison MBs.^54^ This has been attributed this to albumin-shelled MBs having a lower stability *in vivo* compared to phospholipid shells.^57^ This may have important implications where PCD is used to provide real-time feedback control on MB activity by adjusting the ultrasound PNP.^58–60^

Regarding microbubble size, little variation was noted for each sonication target (Fig. 2E, top), and no significant differences were identified in the average HCC among sizes (Fig. 2E, middle). The resulting HCC for each size was as follows: 7.4 ± 2.4, 7.4 ± 1.3 and 7.6 ± 0.6 V2*s for 1 µm, 3 µm and 5 µm MBs, respectively. BCC showed similar trends with no significant differences between the sizes (2.2 ± 0.6, 2.4 ± 0.5 and 2.6 ± 0.1 V2*s for 1 µm, 3 µm and 5 µm MBs, respectively) (Fig. 2E, bottom).

### BBB Opening is Dependent on Microbubble Composition and Number of Sonication Points

Figure 3A shows the region of the mouse brain receiving MB+FUS treatments and BBBO quantification. An initial set of tests was conducted to determine the optimal time to image post-Gd injection. Figure 3B shows the contrast enhancement on T1w MRI for two distinct mechanical indexes: 0.3 (0.3 MPa in situ) and 0.4 (0.4 MPa in situ). Both pressures exhibited a rapid increase within 0 to 10 min, followed by a gradual decline after 15 min. Hence, an imaging time of 14 min was selected.

**Figure 3:**
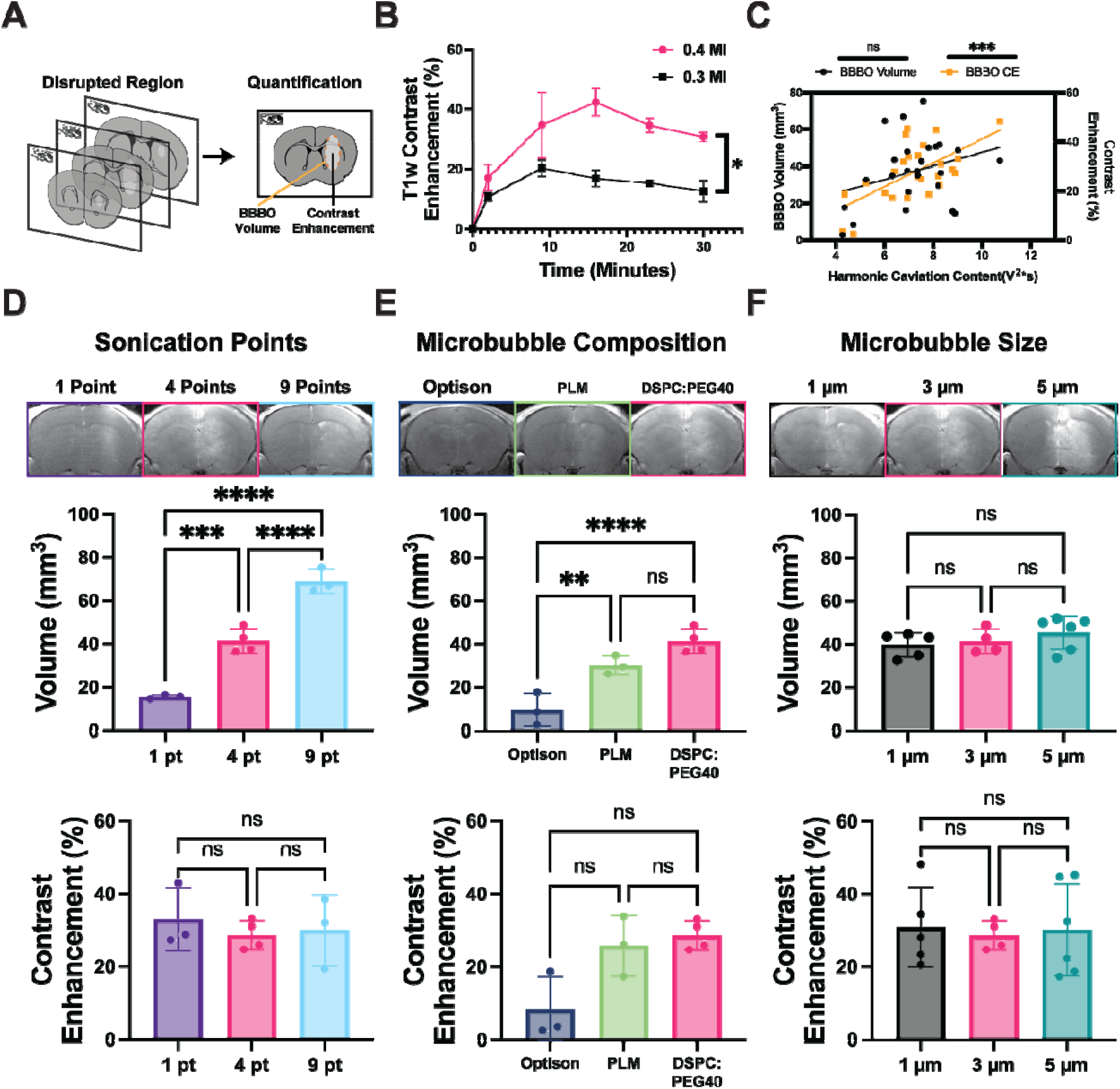
Quantification of BBB Opening via MRI. (A) Cartoon of T1w+CE MRI signal intensit quantification method. (B) T1w+CE contrast enhancement changes over time directly after BBB opening fo two mechanical indices (0.4 and 0.3). (C) Correlation between BBBO volume (black) / BBBO contrast enhancement (orange) and harmonic cavitation content. MRI was quantified for each group and compared between each variation: sonication points (D), composition (E) and size (F). Each panel (D-F) T1w+CE MRI representative images (top) are shown for each group. The total BBB opening volume (middle) and contrast enhancement (bottom) are also shown. Data are repeated for 4 points / 3 um / DSPC:PEG40S to make direct comparisons to other groups. All data are represented as mean ± SD.

Figure 3C shows a correlation analysis between average HCC and the extent of BBBO (volume and contrast enhancement). There was no significant correlation between BBBO volume and average HCC, as expected due to the primary change being the number of sonication points, which showed similar HCC per point. However, a significant linear trend was observed between average HCC and contrast enhancement, likely attributed to the consistency in sonication points.

Figure 3D shows the difference in BBBO between the number of sonications was evident. With an increasing number of sonication points, there was an observable increase in the thickness of vertical streaks on T1w MRI (Fig. 3D, top). This is quantified in BBBO volume, which significantly increased from 15.5 ± 0.9 to 41.3 ± 5.7 then 68.9 ± 5.7 mm^3^ for 1, 4 and 9 points, respectively (Fig. 3D, middle). However, this trend did not hold for contrast enhancement, which showed no significant difference between groups (33.0 ± 8.7, 28.7 ± 4, 30.0 ± 9.8 % for 1, 4 and 9 points, respectively) (Fig. 3D, bottom). Even at a similar PNP (0.4 MPa, in situ) and MVD (10 µL/kg), a wide variety of BBBO volumes was observed, illustrating the importance of MB formulation. As anticipated, the number of sonications increased the BBBO volume (Figure 3D). However, the observed increase in volume did not align directly with the expected volume for the number of points. For instance, given the average volume of 15.5 ± 0.9 mm^3^ for a single point, at 4 points the expected volume would be near 62 mm^3^, yet the observed volume was 41.3 ± 5.7 mm^3^. Similarly, at 9 points 139.5 mm^3^ was anticipated, but the observed volume was only 68.9 ± 5.7 mm3. This discrepancy may be attributed to the potential overlapping of sonication points and skull effects, as evidenced by the drop-off in HCC at the lower right side (Fig. 3D, top). Additionally, a decay in MB concentration within the circulation due to clearance as the FUS system was moved from one sonication point to the next may have caused the decreased BBBO volume.

Microbubble composition influenced BBBO volume, as evident in representative T1w MRI (Fig. 3E, top). Differences were observed between protein-shelled Optison (9.6 ± 7.6 mm^3^) and lipid-shelled PLM (30.2 ± 4.3 mm^3^) or DSPC:PEG40S MBs (41.3 ± 5.7 mm^3^) (Fig. 3E, middle). Although not statistically significant, a similar trend was observed in contrast enhancement, with Optison yielding less BBBO than the other compositions (8.2 ± 9.1, 25.7 ± 8.3 and 28.7 ± 4.0 % for Optison, PLM and DSPC:PEG40S respectively) (Fig. 3E, bottom). Thus, Optison MBs resulted in a significantly smaller BBBO volume and lower contrast enhancement compared to either lipid-shelled MB (PLM or DSPC:PEG40S). Similar to the PCD analysis, prior research has previously observed poorer performance by protein-shelled MBs.^61^ For example, Mcdannold et al. showed similar BBBO at a 5 times higher dose of Optison compared to Definity.^61^ We also show poor Optison stability in rodent blood, which may explain the poor BBBO performance in our study (Supplemental Figure S4).

Microbubble size had minimal influence on both the BBBO volume and contrast enhancement (Fig. 3F). The resulting volumes were as follows: 39.7 ± 5.6, 41.3 ± 5.7 and 45.3 ± 7.7 mm^3^ for 1 µm, 3 µm and 5 µm MBs, respectively (Fig. 3F, middle). Similarly, contrast enhancement showed no significant differences among sizes (30.1 ± 11.0, 28.7 ± 4.0 and 30.1 ± 12.7 % for 1 µm, 3 µm and 5 µm MBs, respectively) (Fig. 3E, bottom). The negligible effect of microbubble size on BBBO extent is consistent with previous findings, where Song et al. showed that when matched at MVD, the extent of BBBO is similar in terms of Evan’s Blue extravasation.^62^

### SIR Measurement by RNA Sequencing Illustrates Varying Levels of Differential Gene Expression

Brain samples were collected 6 h post-sonication from treated regions and subjected to bulk RNA sequencing (Fig. 4A). Differentially expressed genes (DEGs) were analyzed, and volcano plots were generated to illustrate the DEGs in each group (Fig. 4B-D). Figure 4B shows that, as the number of sonication points increased, the number of DEGs also increased, rising from 58 (1 point) to 539 (4 points) to 2184 (9 points). Notably, the percentage of upregulated genes also increased (48, 63 and 71 % for 1, 4 and 9 points, respectively) (Fig. 4B). Previous researchers have described a set of genes heavily associated with SIR, shown in Figure 4E.^22,41^ Each of these sets of differential expressions was compared to the control group. The number of sonication points trended to higher expression of almost all genes except Timp1 and Tlr1, where they showed similar expression in all groups (Fig. 4E, left).

**Figure 4:**
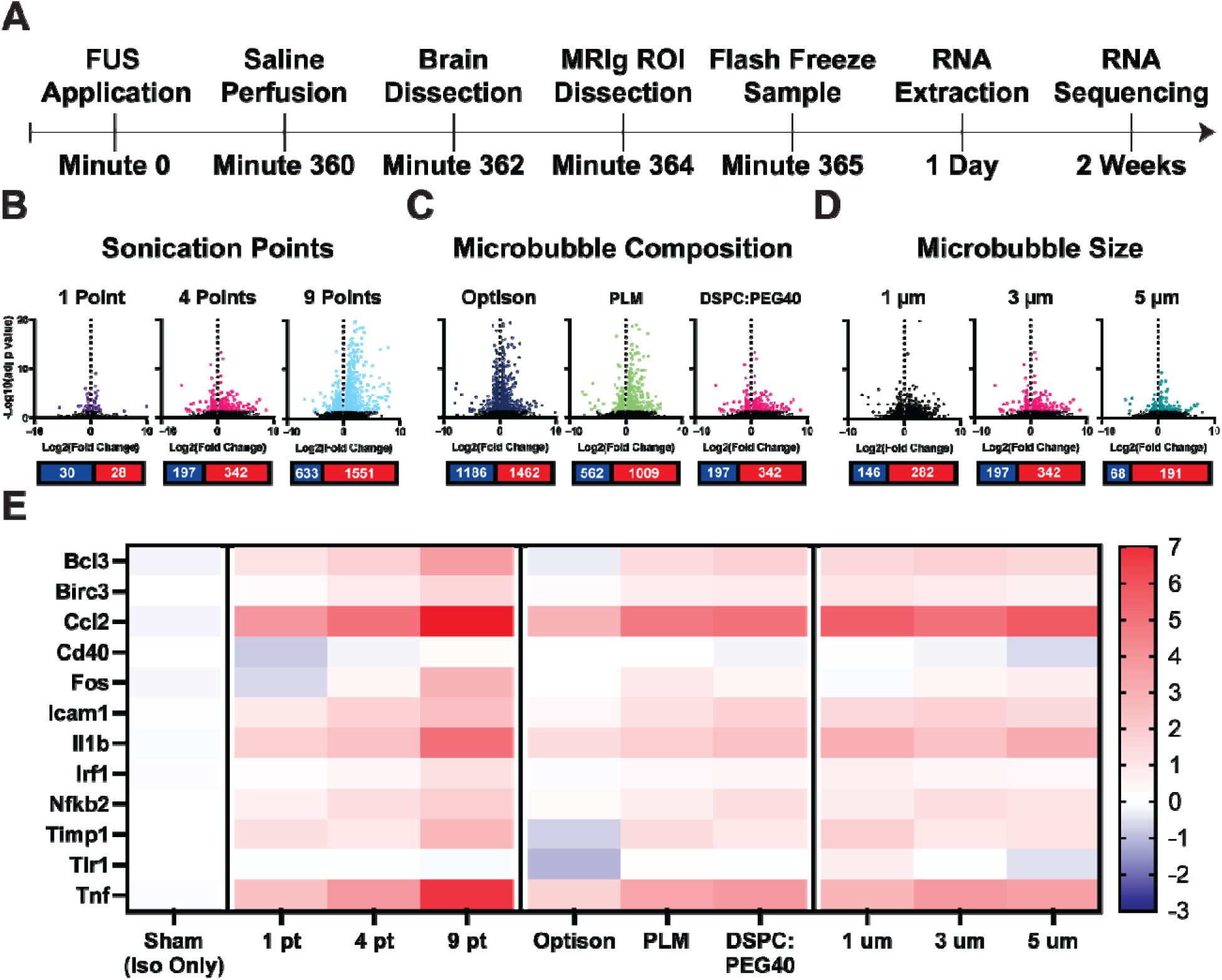
Differential Gene Expression Analysis. (A) Timeline of samples after FUS application until RNA sequencing. DEGs were found for each group: sonication points (B), composition (C) and size (D). The number of DEGs are shown below with upregulated genes shown in red and downregulated genes shown in blue. (E) The fold change for a set of previously described genes associated with SIR. Data was repeated for 4 points / 3 um / DSPC:PEG40S to make direct comparisons to other groups. All data are represented as mean ± SD.

Microbubble composition had diverse effects on the presented DEGs (Fig. 4C). Optison exhibited the highest number of DEGs (2648), although it did not have the highest percentage of upregulated genes (55%). PLM followed with 1571 DEGs, with 64% showing upregulation. DSPC:PEG40S MBs displayed the lowest number of DEGs (539) and the second highest percentage of upregulated genes (63%). Microbubble composition had a dividing line between protein-shelled and lipid-shelled microbubbles, where Optison had lower expression levels in all genes, even showing some downregulation in Tlr1 (Fig. 4E, middle). Both lipid-shelled MBs showed similar expression levels in all genes (Fig. 4E, middle).

Figure 4D shows that MB size demonstrated similar amounts of DEGs, with 408, 539 and 259 for 1, 3 and 5 µm MBs, respectively. Notably, the 5 µm diameter showed a higher percentage of upregulated genes (74%) compared to 1 µm (64%) and 3 µm (63%). However, MB size showed no noticeable trends in terms of gene expression levels (Fig. 4E, right).

A single contralateral control was sent in for RNA sequencing (9 points) to determine the effects on the contralateral hemisphere. DEGs were analyzed similarly to the other groups and showed lower DEGs (70), although still higher than a single sonication point (58) (Fig. 5A). However, the percentage of upregulated genes was lower (44%). Interestingly, when expression levels of SIR-associated genes were compared to the control group, downregulation was observed in six of the twelve genes, with a notable decrease in ccl2 expression (Fig. 5B).

**Figure 5:**
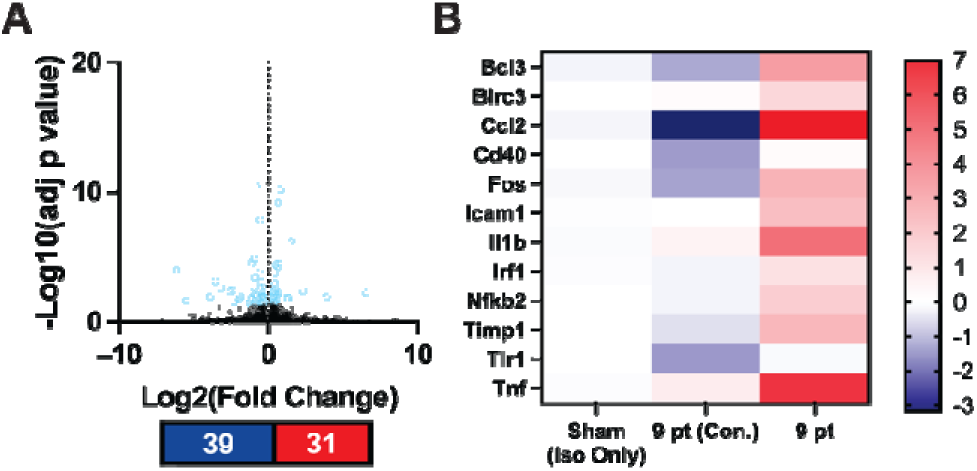
Contralateral Sample Analysis. (A) The contralateral side of the 9-point group was analyzed for DEGs and shown on the volcano plot. The number of DEGs are shown below with upregulated genes shown in red and downregulated genes shown in blue. (B) The fold change for a set of previously described genes associated with SIR. All data are represented as mean ± SD.

Differentially expressed genes provide insights into expression differences between groups. There is a general trend of increased DEGs with an increase in the number of sonication points, extending to a specified gene set associated with SIR (Fig. 4B,E). One intriguing finding is the downregulation of half of the SIR-implicated genes observed in the contralateral side of 9 points (Fig. 5). This underscores the importance of using a single replicate per animal, as multiple samples from the same brain may influence initial inflammatory responses from those regions.^42,63^ The influence of MB composition is multifaceted, with Optison showing more DEGs compared to PLM or DSPC:PEG40S MBs, although expression levels of SIR-specific gene sets showed downregulation or similar expression levels to the control (Fig. 4C,E). This may be explained by the lower *in vivo* stability of Optison compared to the lipid-shelled MBs. Interestingly, MB size did not significantly affect upregulation in the gene set (Fig. 4D,E).

### GSEA Shows Differences in Inflammatory Response Varies Between All Groups

A GSEA was conducted to contextualize the DEGs within biological pathways using a well-established 50 hallmark gene set. Figure 6A shows the GSEA of 38 of the 50 hallmark pathways (the remaining 12 were not significant in any group and therefore not shown). There is a significant increase in normalized enrichment score (NES) as the number of sonication points was increased (Fig. 6A, left). This trend is also evident in the number of significantly upregulated gene sets (12, 23 and 38 for 1, 4 and 9 points, respectively). Of the upregulated gene sets, the two highest NES were found in TNFA signaling via NFKB and inflammatory response. Interestingly, of the 38 upregulated gene sets, the contralateral side of the 9-point group showed significant downregulation in 10 gene sets (including TNFA signaling via NFKB and inflammatory response) and only significant upregulation in 2 gene sets.

**Figure 6:**
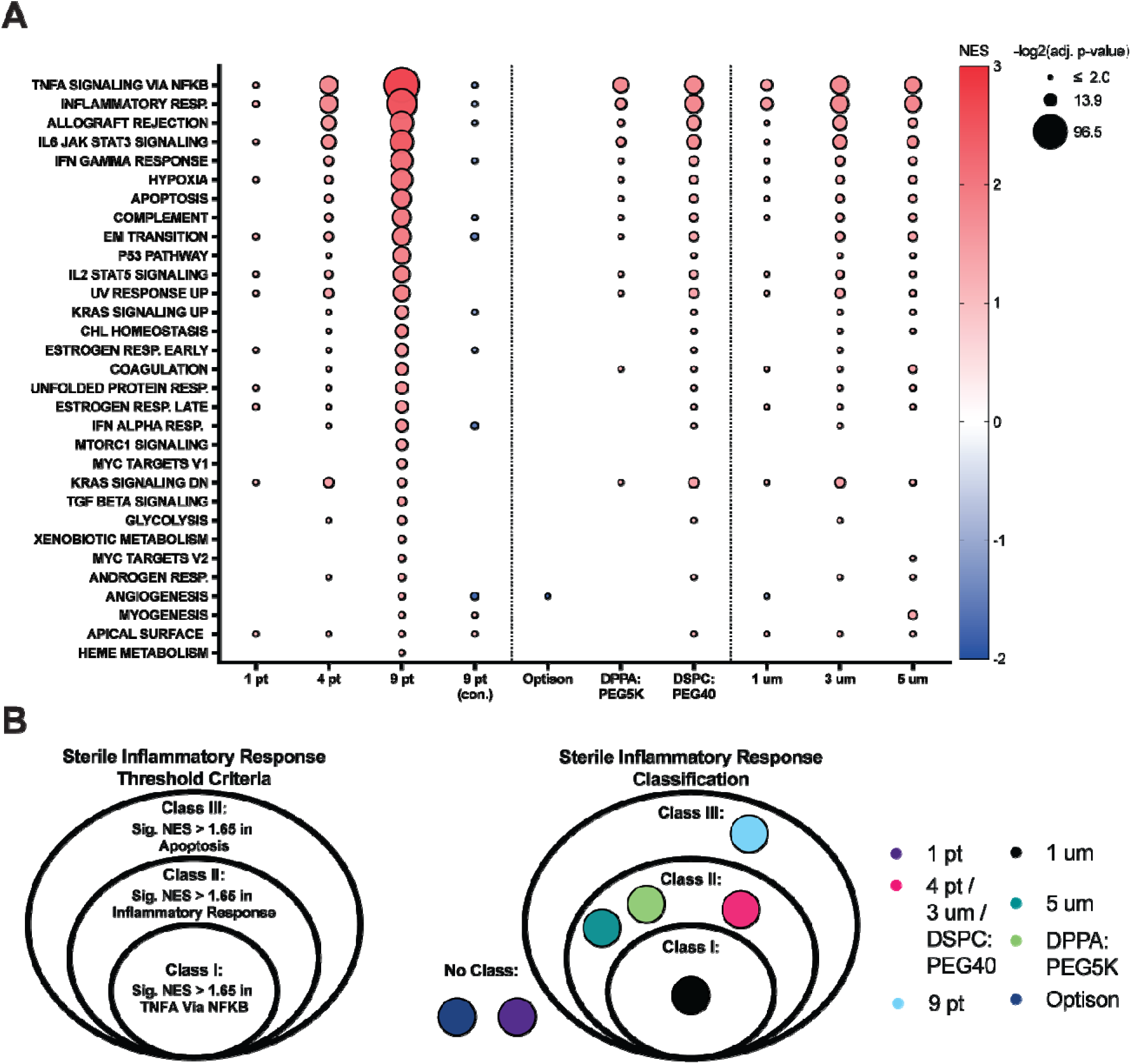
Gene Set Enrichment Analysis. (A) Gene set enrichment analysis was conducted for the 5 hallmark gene sets. The top 30 significant gene sets are plotted. The normalized enrichment score is illustrated via color from blue (downregulated) to red (upregulated). The size of each bubble represents the adjusted p-value. Only significantly normalized enrichment scores are plotted. (B) SIR classifications previously defined by Martinez and Song et al. (left) were used to classify each group used in this study (right) ^23^. Any groups that did not fall within a classification are shown to the left of the classes. Data ar repeated for 4 points / 3 um / DSPC:PEG40S to make direct comparisons to other groups.

The 50-hallmark GSEA provided valuable biological insights into the response observed in each group. The largest range in SIR responses was observed in the number of sonications groups. The substantial increase in SIR magnitude, as explained by NES values from the GSEA and higher classification, illustrates a strong influence of the number of sonication points on the SIR. While many studies have investigated the effects of other parameters on SIR, the variation in the number of sonication points across studies makes direct comparisons challenging.^21–23,43^

The GSEA is also shown for each microbubble composition (Fig. 6A, middle). A clear distinction was observed between the protein-shelled (Optison) and lipid-shelled (PLM and DSPC:PEG40S) MBs. Optison showed a single significant gene set, angiogenesis, which was downregulated. In contrast, PLM and DSPC:PEG40S MBs exhibited similar upregulation of inflammatory gene sets (including TNFA signaling via NFKB and inflammatory response). However, DSPC:PEG40S MBs showed a higher number of significantly upregulated gene sets (23) compared to PLMs (13).

Microbubble composition is another parameter often not controlled between studies, as highlighted in the debate between Kovacs et al.^40,41^ and McMahon and Hynynen.^42,43^ Here, a direct comparison between the three compositions revealed that albumin-shelled MBs (Optison) induced a much smaller SIR, in addition to lower BBBO, at similar PNP and MVD compared to lipid-shelled MBs (PLM or DSPC:PEG40S). This result may be due to the relative biological instability of Optison (Supplementary Fig. S4). Small differences between the two lipid-shell MBs were also observed, indicating a range of responses depending on MB composition. Interestingly, there were no major variations in the type of response between each composition, but rather differences in magnitude.

Overall, each MB size showed similar upregulation in the same gene sets with the same order of increasing NES values. However, MB size exhibited differences in the magnitude of the GSEA (Fig. 6A, right). The smallest size (1 µm diameter) showed fewer upregulated gene sets and had a lower NES for both TNFA signaling via NFKB (1.59) and inflammatory response (1.66). Conversely, 3 and 5 µm MBs had similar NES values for TNFA signaling via NFKB (1.83 and 1.87 for 3 and 5 µm, respectively) and inflammatory response (1.84 and 1.88 for 3 and 5 µm, respectively).

Microbubble size showed small changes in the magnitude of SIR, with smaller MBs exhibiting a smaller magnitude of SIR illustrated by NES and classification. However, this difference was not as pronounced as the effects of the number of sonication points or MB shell type. Interestingly, only minor differences were observed between the 3 and 5 µm MBs. One potential explanation for this result is the larger polydispersity of the largest (5 µm) MBs compared to the other two sizes, leading to a greater variation in DEGs and therefore lower GSEA scores.

Martinez and Song et al. previously described a class system for the magnitude of SIR dependent on the NES found in a GSEA (Fig. 6B, left).^23^ Using this classification system, each group was placed in its respective classification (Fig. 6B, right). Notably, any group that did not fit the lowest criteria was placed off to the side, indicating SIR was not invoked. Two groups, Optison and 1-point sonication with DSPC:PEG40S 3-µm diameter, fit this description, with no significant NES in TNFA signaling via NFKB above 1.65. In Class I, 1 µm MBs were alone indicating a lower SIR than the larger 3 and 5 µm MBs. Class II included three groups (PLM, 3 and 5 µm MBs). Finally, the 9-point sonication exhibited a significant NES in apoptosis (> 1.65), to be placed in the highest class of SIR (Class III).

### The Extent of BBB Opening and Harmonic Cavitation Content are the Strongest Indicators for SIR

Once all data were collected, a correlation matrix was conducted comparing the extent of BBBO (volume and contrast enhancement), PCD values (average HCC/BCC and total HCC/BCC) and the resulting NES of the GSEA (Fig. 7A). A clear correlation was observed between total HCC and BBBO volume (R = 0.92). Additionally, a significant correlation was found between average HCC and BBBO contrast enhancement (R = 0.96). There were no significant correlations between BCC and most other dependent variables. However, there were a few significant correlations between BCC and NES. These include average BCC and allograft rejection (R = −1.0), total BCC and TNFA signaling via NFKB (R = 0.92) or complement (R = 1.0). The distinction between the two is likely due to the consistent nature of each sonication point. If the PNP or MVD changed between groups, there may not have been as strong of a correlation, since previous studies have shown lower correlation significance when either of these two parameters was altered.^60,64,65^ The correlation matrix also included 10 gene sets found in the 50-hallmark gene sets (only those with at least 3 groups with significant NES). Of these, two metrics showed the strongest correlations: BBBO volume had significant positive correlations with 6 groups (R > 0.93), and total HCC illustrated 7 significant positive correlations (R > 0.95).

**Figure 7:**
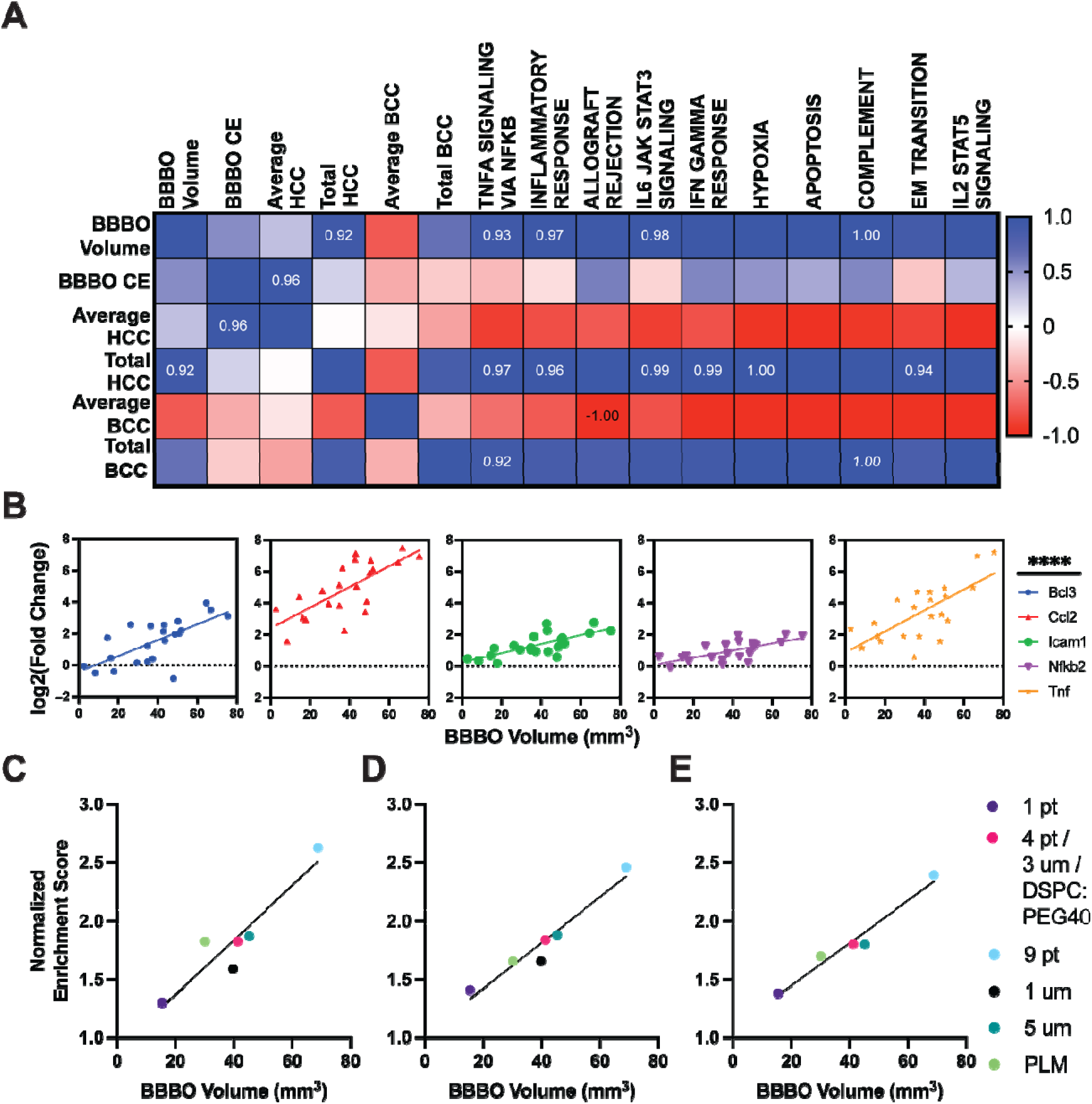
Interplay of Blood-Brain Barrier Opening, Passive Cavitation Detection and Gene Expression. (A) Correlation matrix between dependent variables (BBBO volume, BBBO CE, average HCC, total HCC, average BCC, total BCC) to the top 10 most upregulated gene sets found previously. Significant Pearson R values are shown on the matrix directly. Of the top BBBO volume correlations ones that had at least 5 values were plotted to a linear regression. These included TNFA signaling via NFKB. (B) The fold change for each gene associated with SIR was compared to the resulting BBBO volume. The most significant linear regressions of the gene set are shown. (C), inflammatory response (D) and IL6 jak stat3 signaling (E). Dots are different colors to illustrate the corresponding group.

Of the gene set previously associated in the SIR, particularly by the stimulus of FUS medicated BBBO, five showed significant linear trends with BBBO volume. Figure 7B illustrates each of these five genes (Bcl3, Ccl2, Icam1, Nfkb2 and tnf). The resulting R^2^ values were: 0.47, 0.52, 0.53, 0.46 and .50 respectively. This linear dependence of inflammatory pathways with BBBO has been shown previously either using qPCR^22,66^ or GSEA scores.^23^ However, both have been compared to the contrast enhancement rather than the total volume of the opening.

Three gene sets showed significant upregulation across more than five groups: TNFA signaling via NFKB, inflammatory response and IL6 jak stat3 signaling (Figure 7C-E). When plotted against BBBO volume, a significant linear regression was observed in all gene sets (R2 = 0.87, 0.94, 0.96 for TNFA signaling via NFKB, inflammatory response and IL6 jak stat3, respectively). Interestingly, two minor deviations from the linear regressions were observed. First, PLMs exhibited a higher TNFA signaling via NFKB NES value for the respective BBBO volume (Fig. 7C), although this was not reflected in the other gene sets. Second, smaller (1 µm) MBs demonstrated a lower NES score for TNFA signaling via NFKB (Fig. 7C) and inflammatory response (Fig. 7D), and the effect was insignificant IL6 jak stat3 signaling NES (Fig. 7D). To our knowledge, only one other study has compared microbubble size influence on SIR.^66^ McMahon et al. showed higher relative expression for inflammatory genes in larger microbubbles; however, the use of different microbubble compositions made the influence unclear.^66^

## LIMITATIONS

Despite the valuable insights provided by our study, several limitations should be acknowledged. Firstly, our study focused on bulk RNA sequencing to analyze gene expression changes associated with SIR. While this approach provides valuable information, it may overlook cell-specific responses and dynamic changes in gene expression over time. Incorporating single-cell RNA sequencing and longitudinal studies could provide a more nuanced understanding of the cellular dynamics underlying SIR following BBBO. Additionally, our study primarily utilized mouse models, which may not fully recapitulate the complexities of human physiology and pathology. Translation of our findings to clinical settings warrants careful consideration of species differences and potential confounding factors. Lastly, our study focused on a select set of hallmark gene sets to characterize the inflammatory response following BBBO. While these gene sets provide insights into broad biological processes, they may not fully capture the complexity of SIR. Future studies could explore additional gene sets and pathway analyses to gain a more comprehensive understanding of the molecular mechanisms underlying the SIR. Given these limitations, our study contributes useful insights into the factors influencing BBBO and subsequent inflammatory responses in the brain, laying the groundwork for future research aimed at optimizing drug delivery strategies and mitigating adverse inflammatory reactions.

## CONCLUSION

We investigated the influence of various FUS (number of sonications) and MB (composition and size) parameters on BBBO and subsequent SIR. Our findings underscore the critical role of these parameters in modulating the extent and nature of BBBO and highlight the intricate interplay between microbubble formulation characteristics, sonication parameters and induced inflammatory pathways. We demonstrated that the number of sonication points, as well as microbubble composition, significantly affect BBBO volume and contrast enhancement.

Specifically, increasing the number of sonication points led to an increase in BBBO volume, while microbubble composition exhibited distinct effects on BBBO extent and SIR induction. Notably, lipid-shelled microbubbles demonstrated a higher BBBO volume and a more pronounced inflammatory response compared to protein-shelled microbubbles at a matched MVD. Furthermore, our correlation analysis revealed strong associations between PCD measurements, BBBO parameters and the expression of inflammatory gene sets. Particularly, the correlation between BBBO volume and the upregulation of key inflammatory pathways underscores the critical role of BBBO characteristics and cavitation content in modulating the subsequent sterile inflammatory response. These findings have significant implications for the development of targeted drug delivery strategies and the mitigation of unwanted inflammatory reactions associated with therapeutic interventions targeting the central nervous system.

## ASSOCIATED CONTENT

### Supporting Information

Supplemental Materials (PDF)

## AUTHOR INFORMATION

## Author Contributions

Conceptualization: PM, MB, KHS; Methodology: PM, JS, JD; Investigation: PM, JS, JC; Visualization: PM; Funding acquisition: MB, KHS, PM; Project administration: MB, AG; Supervision: AG, MB; Writing: PM, JS, MB.

## REFERENCES

(1) Hynynen, K.; McDannold, N.; Vykhodtseva, N.; Jolesz, F. A. Noninvasive MR Imaging– Guided Focal Opening of the Blood-Brain Barrier in Rabbits. Radiology 2001, 220 (3), 640–646. 10.1148/radiol.2202001804.

(2) Martinez, P.; Nault, G.; Steiner, J.; Wempe, M. F.; Pierce, A.; Brunt, B.; Slade, M.; Song, J. J.; Mongin, A.; Song, K.-H.; Ellens, N.; Serkova, N.; Green, A. L.; Borden, M. MRI-Guided Focused Ultrasound Blood–Brain Barrier Opening Increases Drug Delivery and Efficacy in a Diffuse Midline Glioma Mouse Model. Neuro-Oncology Advances 2023, 5 (1), vdad111. 10.1093/noajnl/vdad111.

(3) Alli, S.; Figueiredo, C. A.; Golbourn, B.; Sabha, N.; Wu, M. Y.; Bondoc, A.; Luck, A.; Coluccia, D.; Maslink, C.; Smith, C.; Wurdak, H.; Hynynen, K.; O’Reilly, M.; Rutka, J. T. Brainstem Blood Brain Barrier Disruption Using Focused Ultrasound: A Demonstration of Feasibility and Enhanced Doxorubicin Delivery. Journal of Controlled Release 2018, 281, 29–41. 10.1016/j.jconrel.2018.05.005.

(4) Brighi, C.; Salimova, E.; de Veer, M.; Puttick, S.; Egan, G. Translation of Focused Ultrasound for Blood-Brain Barrier Opening in Glioma. Journal of Controlled Release 2022, 345, 443–463. 10.1016/j.jconrel.2022.03.035.

(5) Aryal, M.; Park, J.; Vykhodtseva, N.; Zhang, Y.-Z.; McDannold, N. Enhancement in Blood-Tumor Barrier Permeability and Delivery of Liposomal Doxorubicin Using Focused Ultrasound and Microbubbles: Evaluation during Tumor Progression in a Rat Glioma Model. Phys. Med. Biol. 2015, 60 (6), 2511–2527. 10.1088/0031-9155/60/6/2511.

(6) Ye, D.; Yuan, J.; Yue, Y.; Rubin, J. B.; Chen, H. Focused Ultrasound-Enhanced Delivery of Intranasally Administered Anti-Programmed Cell Death-Ligand 1 Antibody to an Intracranial Murine Glioma Model. Pharmaceutics 2021, 13 (2), 190. 10.3390/pharmaceutics13020190.

(7) Carpentier, A.; Canney, M.; Vignot, A.; Reina, V.; Beccaria, K.; Horodyckid, C.; Karachi, C.; Leclercq, D.; Lafon, C.; Chapelon, J.-Y. Clinical Trial of Blood-Brain Barrier Disruption by Pulsed Ultrasound. Science translational medicine 2016, 8 (343), 343re2–343re2.

(8) Chen, K.-T.; Lin, Y.-J.; Chai, W.-Y.; Lin, C.-J.; Chen, P.-Y.; Huang, C.-Y.; Kuo, J. S.; Liu, H.-L.; Wei, K.-C. Neuronavigation-Guided Focused Ultrasound (NaviFUS) for Transcranial Blood-Brain Barrier Opening in Recurrent Glioblastoma Patients: Clinical Trial Protocol. Ann Transl Med 2020, 8 (11), 673–673. 10.21037/atm-20-344.

(9) Mainprize, T.; Lipsman, N.; Huang, Y.; Meng, Y.; Bethune, A.; Ironside, S.; Heyn, C.; Alkins, R.; Trudeau, M.; Sahgal, A. Blood-Brain Barrier Opening in Primary Brain Tumors with Non-Invasive MR-Guided Focused Ultrasound: A Clinical Safety and Feasibility Study. Scientific reports 2019, 9 (1), 321.

(10) Ferrara, K.; Pollard, R.; Borden, M. Ultrasound Microbubble Contrast Agents: Fundamentals and Application to Gene and Drug Delivery. Annu. Rev. Biomed. Eng. 2007, 9 (1), 415–447. 10.1146/annurev.bioeng.8.061505.095852.

(11) Song, K.-H.; Harvey, B. K.; Borden, M. A. State-of-the-Art of Microbubble-Assisted Blood-Brain Barrier Disruption. Theranostics 2018, 8 (16), 4393–4408. 10.7150/thno.26869.

(12) Navarro-Becerra, J. A.; Song, K.-H.; Martinez, P.; Borden, M. A. Microbubble Size and Dose Effects on Pharmacokinetics. ACS Biomater. Sci. Eng. 2022, 8 (4), 1686–1695. 10.1021/acsbiomaterials.2c00043.

(13) Garg, S.; Thomas, A. A.; Borden, M. A. The Effect of Lipid Monolayer In-Plane Rigidity on in Vivo Microbubble Circulation Persistence. Biomaterials 2013, 34 (28), 6862–6870. 10.1016/j.biomaterials.2013.05.053.

(14) Navarro-Becerra, J. A.; Castillo, J. I.; Borden, M. A. Effect of Poly(Ethylene Glycol) Configuration on Microbubble Pharmacokinetics. ACS Biomater Sci Eng 2024. 10.1021/acsbiomaterials.3c01764.

(15) Bader, K. B.; Holland, C. K. Gauging the Likelihood of Stable Cavitation from Ultrasound Contrast Agents. Phys. Med. Biol. 2013, 58 (1), 127–144. 10.1088/0031-9155/58/1/127.

(16) Stride, E. P.; Coussios, C. C. Cavitation and Contrast: The Use of Bubbles in Ultrasound Imaging and Therapy. Proc Inst Mech Eng H 2010, 224 (2), 171–191. 10.1243/09544119JEIM622.

(17) Bezer, J. H.; Prentice, P.; Kee Chang, W. L.; Morse, S. V.; Christensen-Jeffries, K.; Rowlands, C. J.; Kozlov, A. S.; Choi, J. J. Microbubble Dynamics in Brain Microvessels. October 28, 2023. 10.1101/2023.10.25.563941.

(18) Collis, J.; Manasseh, R.; Liovic, P.; Tho, P.; Ooi, A.; Petkovic-Duran, K.; Zhu, Y. Cavitation Microstreaming and Stress Fields Created by Microbubbles. Ultrasonics 2010, 50 (2), 273–279. 10.1016/j.ultras.2009.10.002.

(19) Khokhlova, T. D.; Haider, Y.; Hwang, J. H. Therapeutic Potential of Ultrasound Microbubbles in Gastrointestinal Oncology: Recent Advances and Future Prospects. Therap Adv Gastroenterol 2015, 8 (6), 384–394. 10.1177/1756283X15592584.

(20) Todd, N.; Angolano, C.; Ferran, C.; Devor, A.; Borsook, D.; McDannold, N. Secondary Effects on Brain Physiology Caused by Focused Ultrasound-Mediated Disruption of the Blood–Brain Barrier. Journal of Controlled Release 2020, 324, 450–459. 10.1016/j.jconrel.2020.05.040.

(21) Kovacs, Z. I.; Kim, S.; Jikaria, N.; Qureshi, F.; Milo, B.; Lewis, B. K.; Bresler, M.; Burks, S. R.; Frank, J. A. Disrupting the Blood–Brain Barrier by Focused Ultrasound Induces Sterile Inflammation. Proc. Natl. Acad. Sci. U.S.A. 2017, 114 (1). 10.1073/pnas.1614777114.

(22) McMahon, D.; Hynynen, K. Acute Inflammatory Response Following Increased Blood-Brain Barrier Permeability Induced by Focused Ultrasound Is Dependent on Microbubble Dose. Theranostics 2017, 7 (16), 3989–4000. 10.7150/thno.21630.

(23) Martinez, P.; Song, J. J.; Garay, F. G.; Song, K.-H.; Mufford, T.; Steiner, J.; DeSisto, J.; Ellens, N.; Serkova, N. J.; Green, A. L.; Borden, M. Comprehensive Assessment of Blood-Brain Barrier Opening and Sterile Inflammatory Response: Unraveling the Therapeutic Window; preprint; Bioengineering, 2023. 10.1101/2023.10.23.563613.

(24) Matzinger, P. Tolerance, Danger, and the Extended Family. Annu. Rev. Immunol. 1994, 12 (1), 991–1045. 10.1146/annurev.iy.12.040194.005015.

(25) Rock, K. L.; Latz, E.; Ontiveros, F.; Kono, H. The Sterile Inflammatory Response. Annu. Rev. Immunol. 2010, 28 (1), 321–342. 10.1146/annurev-immunol-030409-101311.

(26) Zindel, J.; Kubes, P. DAMPs, PAMPs, and LAMPs in Immunity and Sterile Inflammation. Annu. Rev. Pathol. Mech. Dis. 2020, 15 (1), 493–518. 10.1146/annurev-pathmechdis-012419-032847.

(27) Keyel, P. A. How Is Inflammation Initiated? Individual Influences of IL-1, IL-18 and HMGB1. Cytokine 2014, 69 (1), 136–145. 10.1016/j.cyto.2014.03.007.

(28) Chen, Y.; Yousaf, M. N.; Mehal, W. Z. Role of Sterile Inflammation in Fatty Liver Diseases. Liver Research 2018, 2 (1), 21–29. 10.1016/j.livres.2018.02.003.

(29) Ratajczak, M. Z.; Pedziwiatr, D.; Cymer, M.; Kucia, M.; Kucharska-Mazur, J.; Samochowiec, J. Sterile Inflammation of Brain, Due to Activation of Innate Immunity, as a Culprit in Psychiatric Disorders. Front. Psychiatry 2018, 9, 60. 10.3389/fpsyt.2018.00060.

(30) Otani, K.; Shichita, T. Cerebral Sterile Inflammation in Neurodegenerative Diseases. Inflamm Regener 2020, 40 (1), 28. 10.1186/s41232-020-00137-4.

(31) Serkova, N. J. Nanoparticle-Based Magnetic Resonance Imaging on Tumor-Associated Macrophages and Inflammation. Front. Immunol. 2017, 8, 590. 10.3389/fimmu.2017.00590.

(32) Ji, R.; Karakatsani, M. E.; Burgess, M.; Smith, M.; Murillo, M. F.; Konofagou, E. E. Cavitation-Modulated Inflammatory Response Following Focused Ultrasound Blood-Brain Barrier Opening. Journal of Controlled Release 2021, 337, 458–471. 10.1016/j.jconrel.2021.07.042.

(33) Choi, J. J.; Selert, K.; Gao, Z.; Samiotaki, G.; Baseri, B.; Konofagou, E. E. Noninvasive and Localized Blood—Brain Barrier Disruption Using Focused Ultrasound Can Be Achieved at Short Pulse Lengths and Low Pulse Repetition Frequencies. J Cereb Blood Flow Metab 2011, 31 (2), 725–737. 10.1038/jcbfm.2010.155.

(34) Fan, Z.; Chen, D.; Deng, C. X. Characterization of the Dynamic Activities of a Population of Microbubbles Driven by Pulsed Ultrasound Exposure in Sonoporation. Ultrasound in medicine & biology 2014, 40 (6), 1260–1272.

(35) Kovacs, Z. I.; Tu, T.-W.; Sundby, M.; Qureshi, F.; Lewis, B. K.; Jikaria, N.; Burks, S. R.; Frank, J. A. MRI and Histological Evaluation of Pulsed Focused Ultrasound and Microbubbles Treatment Effects in the Brain. Theranostics 2018, 8 (17), 4837–4855. 10.7150/thno.24512.

(36) Lim Kee Chang, W.; Chan, T. G.; Raguseo, F.; Mishra, A.; Chattenton, D.; De Rosales, R. T. M.; Long, N. J.; Morse, S. V. Rapid Short-Pulses of Focused Ultrasound and Microbubbles Deliver a Range of Agent Sizes to the Brain. Sci Rep 2023, 13 (1), 6963. 10.1038/s41598-023-33671-5.

(37) Chopra, R.; Vykhodtseva, N.; Hynynen, K. Influence of Exposure Time and Pressure Amplitude on Blood−Brain-Barrier Opening Using Transcranial Ultrasound Exposures. ACS Chem. Neurosci. 2010, 1 (5), 391–398. 10.1021/cn9000445.

(38) Goertz, D. E.; de Jong, N.; van der Steen, A. F. W. Attenuation and Size Distribution Measurements of Definity^TM^ and Manipulated Definity^TM^ Populations. Ultrasound in Medicine & Biology 2007, 33 (9), 1376–1388. 10.1016/j.ultrasmedbio.2007.03.009.

(39) Bokor, D.; Chambers, J. B.; Rees, P. J.; Mant, T. G.; Luzzani, F.; Spinazzi, A. Clinical Safety of SonoVue^TM^, a New Contrast Agent for Ultrasound Imaging, in Healthy Volunteers and in Patients with Chronic Obstructive Pulmonary Disease. Investigative radiology 2001, 36 (2), 104–109.

(40) Kovacs, Z. I.; Burks, S. R.; Frank, J. A. Focused Ultrasound with Microbubbles Induces Sterile Inflammatory Response Proportional to the Blood Brain Barrier Opening: Attention to Experimental Conditions. Theranostics 2018, 8 (8), 2245–2248. 10.7150/thno.24181.

(41) Kovacs, Z. I.; Kim, S.; Jikaria, N.; Qureshi, F.; Milo, B.; Lewis, B. K.; Bresler, M.; Burks, S. R.; Frank, J. A. Disrupting the Blood–Brain Barrier by Focused Ultrasound Induces Sterile Inflammation. Proc. Natl. Acad. Sci. U.S.A. 2017, 114 (1). 10.1073/pnas.1614777114.

(42) McMahon, D.; Hynynen, K. Acute Inflammatory Response Following Increased Blood-Brain Barrier Permeability Induced by Focused Ultrasound Is Dependent on Microbubble Dose. Theranostics 2017, 7 (16), 3989–4000. 10.7150/thno.21630.

(43) McMahon, D.; Hynynen, K. Reply to Kovacs et al.: Concerning Acute Inflammatory Response Following Focused Ultrasound and Microbubbles in the Brain. Theranostics 2018, 8 (8), 2249–2250. 10.7150/thno.25468.

(44) Feshitan, J. A.; Chen, C. C.; Kwan, J. J.; Borden, M. A. Microbubble Size Isolation by Differential Centrifugation. Journal of Colloid and Interface Science 2009, 329 (2), 316–324. 10.1016/j.jcis.2008.09.066.

(45) Kwan, J. J.; Borden, M. A. Microbubble Dissolution in a Multigas Environment. Langmuir 2010, 26 (9), 6542–6548. 10.1021/la904088p.

(46) Kwan, J. J.; Borden, M. A. Lipid Monolayer Dilatational Mechanics during Microbubble Gas Exchange. Soft Matter 2012, 8 (17), 4756–4766. 10.1039/C2SM07437K.

(47) Koza, L. A.; Pena, C.; Russell, M.; Smith, A. C.; Molnar, J.; Devine, M.; Serkova, N. J.; Linseman, D. A. Immunocal® Limits Gliosis in Mouse Models of Repetitive Mild-Moderate Traumatic Brain Injury. Brain Research 2023, 1808, 148338. 10.1016/j.brainres.2023.148338.

(48) Martinez, P.; Bottenus, N.; Borden, M. Cavitation Characterization of Size-Isolated Microbubbles in a Vessel Phantom Using Focused Ultrasound. Pharmaceutics 2022, 14 (9), 1925. 10.3390/pharmaceutics14091925.

(49) Love, M. I.; Huber, W.; Anders, S. Moderated Estimation of Fold Change and Dispersion for RNA-Seq Data with DESeq2. Genome Biol 2014, 15 (12), 550. 10.1186/s13059-014-0550-8.

(50) Korotkevich, G.; Sukhov, V.; Budin, N.; Shpak, B.; Artyomov, M. N.; Sergushichev, A. Fast Gene Set Enrichment Analysis; preprint; Bioinformatics, 2016. 10.1101/060012.

(51) Liberzon, A.; Birger, C.; Thorvaldsdóttir, H.; Ghandi, M.; Mesirov, J. P.; Tamayo, P. The Molecular Signatures Database Hallmark Gene Set Collection. Cell Systems 2015, 1 (6), 417–425. 10.1016/j.cels.2015.12.004.

(52) Subramanian, A.; Tamayo, P.; Mootha, V. K.; Mukherjee, S.; Ebert, B. L.; Gillette, M. A.; Paulovich, A.; Pomeroy, S. L.; Golub, T. R.; Lander, E. S.; Mesirov, J. P. Gene Set Enrichment Analysis: A Knowledge-Based Approach for Interpreting Genome-Wide Expression Profiles. Proc. Natl. Acad. Sci. U.S.A. 2005, 102 (43), 15545–15550. 10.1073/pnas.0506580102.

(53) Benjamini, Y.; Hochberg, Y. Controlling the False Discovery Rate: A Practical and Powerful Approach to Multiple Testing. Journal of the Royal Statistical Society. Series B (Methodological) 57 (1 (1995)), 289–300.

(54) Bing, C.; Hong, Y.; Hernandez, C.; Rich, M.; Cheng, B.; Munaweera, I.; Szczepanski, D.; Xi, Y.; Bolding, M.; Exner, A.; Chopra, R. Characterization of Different Bubble Formulations for Blood-Brain Barrier Opening Using a Focused Ultrasound System with Acoustic Feedback Control. Sci Rep 2018, 8 (1), 7986. 10.1038/s41598-018-26330-7.

(55) Keller, S. B.; Sheeran, P. S.; Averkiou, M. A. Cavitation Therapy Monitoring of Commercial Microbubbles With a Clinical Scanner. *IEEE Trans. Ultrason., Ferroelect.*, Freq. Contr. 2021, 68 (4), 1144–1154. 10.1109/TUFFC.2020.3034532.

(56) Newsome, I. G.; Kierski, T. M.; Dayton, P. A. Assessment of the Superharmonic Response of Microbubble Contrast Agents for Acoustic Angiography as a Function of Microbubble Parameters. Ultrasound in Medicine & Biology 2019, 45 (9), 2515–2524. 10.1016/j.ultrasmedbio.2019.04.027.

(57) Chomas, J. E.; Dayton, P.; Allen, J.; Morgan, K.; Ferrara, K. W. Mechanisms of Contrast Agent Destruction. IEEE Transactions on Ultrasonics, Ferroelectrics, and Frequency Control 2001, 48 (1), 232–248. 10.1109/58.896136.

(58) Masiero, M.; Boulos, P.; Crake, C.; Rowe, C.; Coviello, C. M. Ultrasound-Induced Cavitation and Passive Acoustic Mapping: SonoTran Platform Performance and Short-Term Safety in a Large-Animal Model. Ultrasound in Medicine & Biology 2022, 48 (8), 1681–1690. 10.1016/j.ultrasmedbio.2022.03.010.

(59) Wu, Q.; Gray, M.; Smith, C.; Bau, L.; Coussios, C.; Stride, E. P. Correlating High-Speed Optical Imaging and Passive Acoustic Mapping of Cavitation Dynamics. The Journal of the Acoustical Society of America 2022, 151 (4), A174–A174. 10.1121/10.0011017.

(60) Xu, S.; Ye, D.; Wan, L.; Shentu, Y.; Yue, Y.; Wan, M.; Chen, H. Correlation Between Brain Tissue Damage and Inertial Cavitation Dose Quantified Using Passive Cavitation Imaging. Ultrasound in Medicine & Biology 2019, 45 (10), 2758–2766. 10.1016/j.ultrasmedbio.2019.07.004.

(61) McDannold, N.; Vykhodtseva, N.; Hynynen, K. Use of Ultrasound Pulses Combined with Definity for Targeted Blood-Brain Barrier Disruption: A Feasibility Study. Ultrasound in Medicine & Biology 2007, 33 (4), 584–590. 10.1016/j.ultrasmedbio.2006.10.004.

(62) Song, K.-H.; Fan, A. C.; Hinkle, J. J.; Newman, J.; Borden, M. A.; Harvey, B. K. Microbubble Gas Volume: A Unifying Dose Parameter in Blood-Brain Barrier Opening by Focused Ultrasound. Theranostics 2017, 7 (1), 144–152. 10.7150/thno.15987.

(63) McMahon, D.; Bendayan, R.; Hynynen, K. Acute Effects of Focused Ultrasound-Induced Increases in Blood-Brain Barrier Permeability on Rat Microvascular Transcriptome. Sci Rep 2017, 7 (1), 45657. 10.1038/srep45657.

(64) O’Reilly, M. A.; Hynynen, K. Blood-Brain Barrier: Real-Time Feedback-Controlled Focused Ultrasound Disruption by Using an Acoustic Emissions-Based Controller. Radiology 2012, 263 (1), 96–106. 10.1148/radiol.11111417.

(65) McMahon, D.; O’Reilly, M. A.; Hynynen, K. Therapeutic Agent Delivery Across the Blood–Brain Barrier Using Focused Ultrasound. Annu. Rev. Biomed. Eng. 2021, 23 (1), 89–113. 10.1146/annurev-bioeng-062117-121238.

(66) McMahon, D.; Lassus, A.; Gaud, E.; Jeannot, V.; Hynynen, K. Microbubble Formulation Influences Inflammatory Response to Focused Ultrasound Exposure in the Brain. Sci Rep 2020, 10 (1), 21534. 10.1038/s41598-020-78657-9.

