## Supplemental Materials for "Effect of Microbubble Size, Composition and Multiple Sonication Points on Sterile Inflammatory Response in Focused Ultrasound-Mediated Blood-Brain Barrier Opening"

**The PDF file includes:**

Fig. S1: **Size isolation procedure for three microbubble sizes.**
Fig. S2: **Volume-weighted size distributions for all microbubble populations.**
Fig. S3: **Skull shape surrounding target region.**
Fig. S4: **Optison Stability in vivo and in vitro.**
Fig. S5: **Microbubble Stability.**

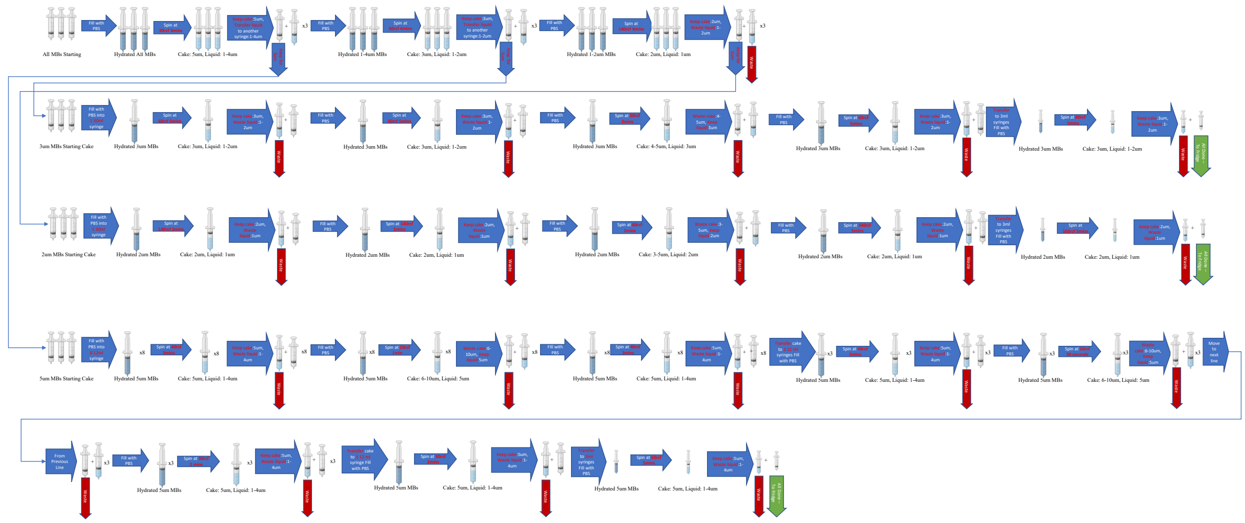

**Figure S1: Size isolation procedure for three microbubble sizes.**

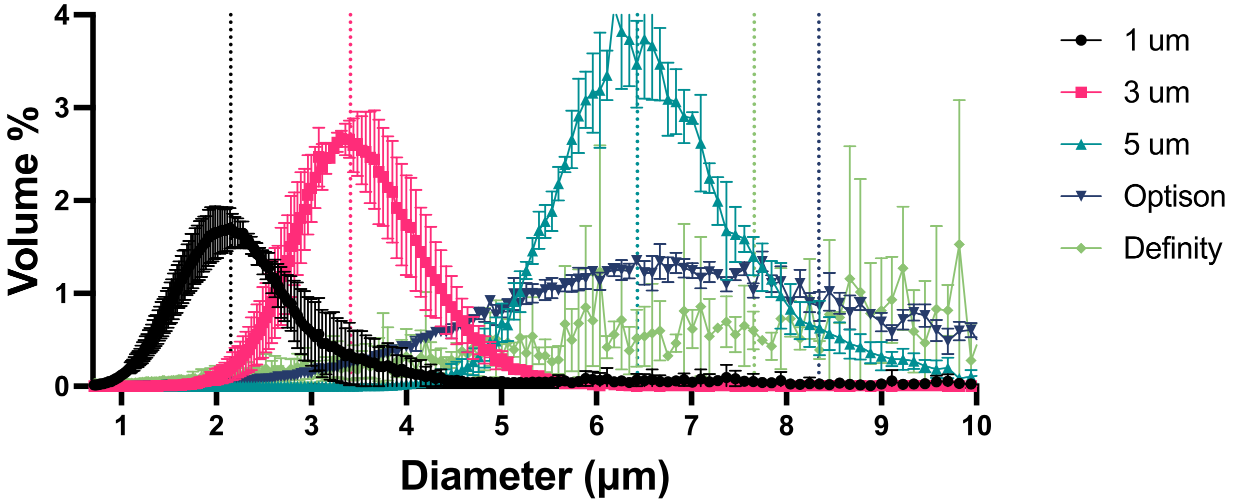


**Figure S2: Volume-weighted size distributions for all microbubble populations.**

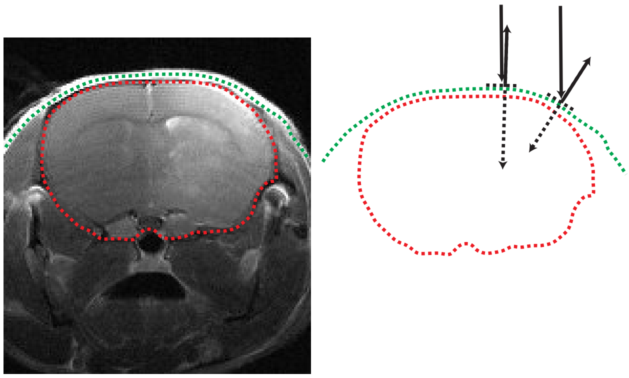

**Figure S7.3: Skull shape surrounding target region. The skull (green) and brain (red) are outlined. Ultrasound is given at a vertical angle from the RK50 system. At the center of the brain, the skull shows little disturbance to vertical FUS application. At regions near the edge, the skull has a larger angle difference to the vertical ultrasound wave.**

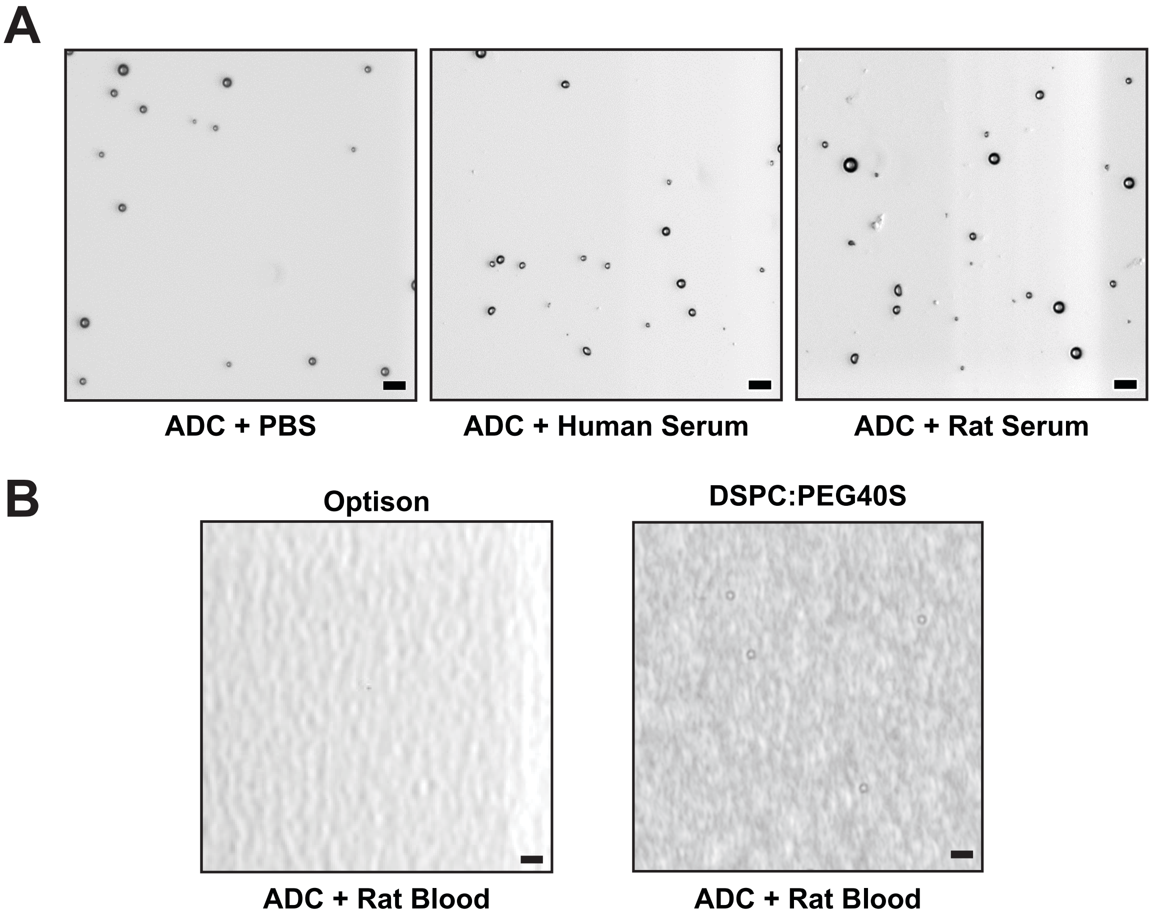


**Supplemental Figure 4: Optison Stability *in vivo* and *in vitro*. (A) *In vitro* testing of Optison, where Optison was mixed and imaged with anticoagulant (ADC) + PBS (left), ADC + human serum (middle left) and ADC + rat serum (middle right). (B) In vivo testing of Optison (left) and DSPC:PEG40S (right) MBs samples were injected into a rat and blood was collected after 2 minutes. Brightfield microscopy is shown for all samples. The scale bar is 10 µm.**

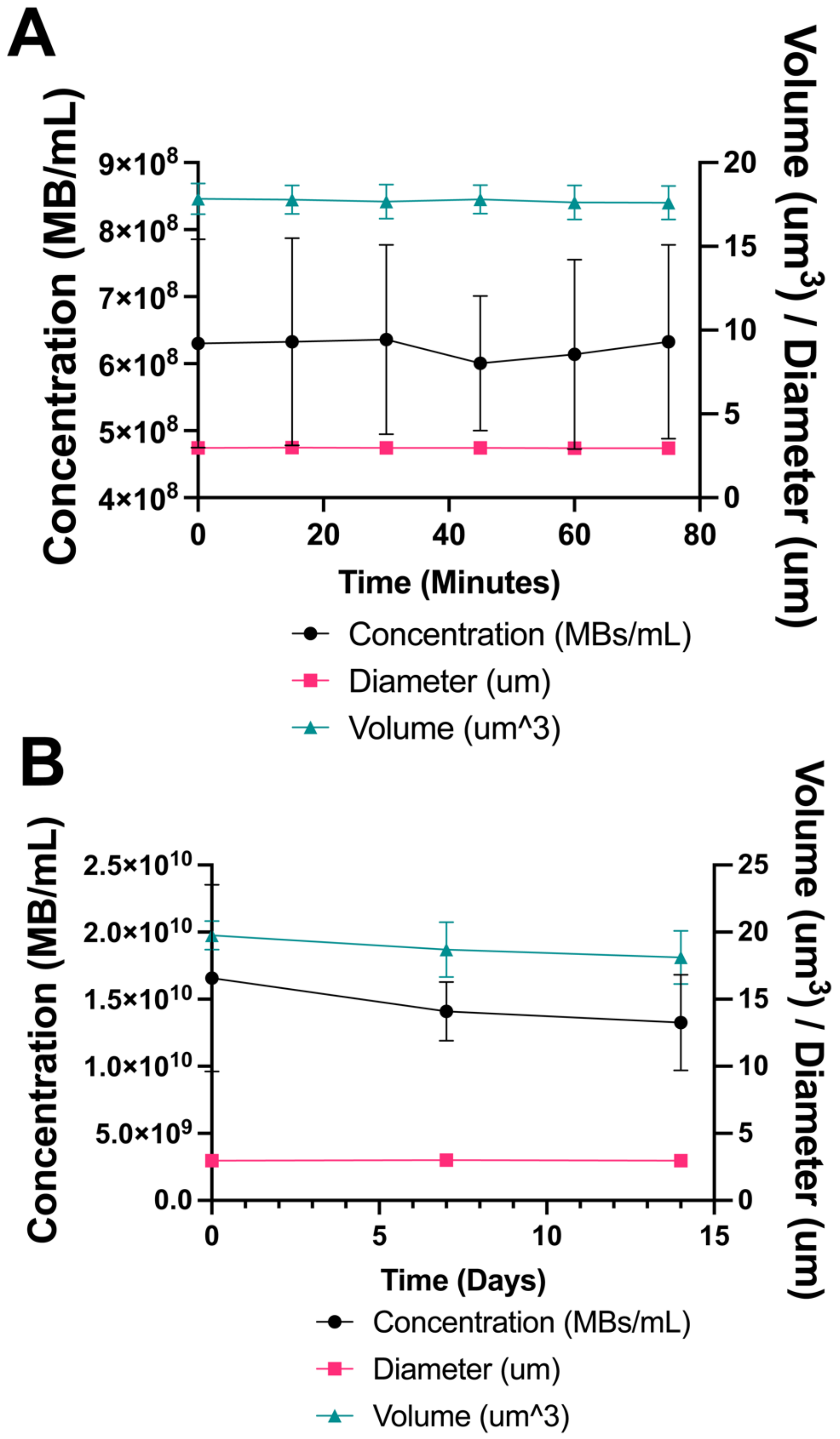


**Supplementary Figure 5. Microbubble Stability.** (A) Measurements of microbubbles after 1 hour at diluted concentrations used during injection (~5E8MBs/mL). (B) The plot shows measurements of microbubbles after 2 weeks of storage at 4C. Data is shown as mean ± SD (*n = 3*).
